# Reproducible Community States Resolve the Nasopharyngeal Diversity Paradox

**DOI:** 10.1101/2025.09.06.673381

**Authors:** Kuncheng Song, Hayden N Brochu, Monica L Bustos, Qimin Zhang, Martia J Thomas, Ayla B Harris, Crystal R Icenhour, Stanley Letovsky, Lakshmanan K Iyer

## Abstract

The nasopharyngeal microbiome is critical for respiratory health, yet its compositional architecture remains largely uncharacterized, underlying conflicting reports of increased, decreased, or unchanged diversity during infection. Here, we demonstrate that these contradictions arise from a fundamentally overlooked factor: the nasopharynx is organized into six reproducible nasopharyngeal community state types (NPCSTs) that are associated with intrinsic diversity baselines independent of disease status. By uniformly reprocessing 7,790 16S rRNA gene sequencing samples from 28 studies, we show that NPCSTs explain 52% of community variance, four-fold more than study effects, and that disease-diversity associations are attenuated by over 92% after NPCST adjustment. To identify genuine disease markers, we applied NPCST-aware differential abundance testing to SARS-CoV-2 as a case study. Leave-one-study-out (LOSO) cross-validation across 10 cohorts confirmed 13 reproducible taxa with opposing shifts: obligate anaerobes typically studied in other body compartments were enriched, while resident hub genera were depleted with high directional consistency across LOSO iterations. Co-occurrence network analysis independently validated this pattern, identifying the depleted taxa as structural backbone hubs across all NPCSTs. To bridge microbiome ecology and clinical application, we developed the Nasopharyngeal Microbiome Health Index (NMHI), a continuous community-level wellness score achieving AUC of 0.90 and 0.92 in internal and external validations, respectively, with robust performance across all six NPCSTs (AUC: 0.848-0.953). An independent clinical cohort of 147 specimens confirmed that these NPCST and NMHI characteristics are reproducible. Unlike binary classifiers, NMHI quantifies nasopharyngeal health along a spectrum, establishing NPCST-aware analysis as a foundation for precision respiratory microbiome research.

## Introduction

The nasopharynx serves as a critical reservoir of microbial colonization, where its anatomically protected mucosa harbors diverse microbes [1]. Established as the preferred reference site for respiratory diagnostics [2, 3], nasopharyngeal sampling underpins PCR-based panels targeting dozens of pathogens [4, 5], with millions of nasopharyngeal swabs collected annually in the United States during the COVID-19 pandemic alone [6–8]. However, as a distinct microbial ecosystem at the interface between the external environment and lower airway, the core nasopharyngeal microbiome community structure has not been systematically characterized. This knowledge gap became critically apparent with the COVID-19 pandemic, when conflicting studies reported increased [9, 10], decreased [11–14], or no alterations [15–17] in diversity between SARS-CoV-2 and control samples.

These contradictions reflect four unresolved methodological challenges. First, contamination from environmental and reagent-derived sources is particularly problematic in low-biomass nasopharyngeal samples [18–20], and existing decontamination methods (decontam [21], micRoclean [22], and CleanSeqU [23]) require negative controls that are unavailable in most published datasets. Second, attempts to define distinct nasopharyngeal community structures have yielded study-specific clusters ranging from 4 to 13 [16, 24–34], with no framework demonstrating reproducibility across independent cohorts. Third, no large-scale comprehensive study of the nasopharyngeal microbiome exists, limiting the field to isolated cohorts. These cohorts’ differential abundance analyses yield study-specific biomarker taxa that lack external validation, while machine learning classifiers targeting individual genera achieve only modest performance (AUC 0.59-0.82) [27, 30, 35–37]. Fourth, clinical translation lacks tools for quantifying overall nasopharyngeal microbiome health. This gap is compounded by poly-microbial colonization, because most respiratory diagnostic and research frameworks target a single pathogen and rarely capture the coexistence of multiple pathogenic and opportunistic agents in one sample.

We hypothesized that these four limitations stem from a fundamentally overlooked factor. The genuine nasopharyngeal microbiome is organized into distinct nasopharyngeal community state types (NPCSTs) that are consistent with intrinsic baselines, and failure to account for this unique architecture confounds both diversity comparisons and differential abundance testing across studies. Under this framework, conflicting findings arise not from disease effects but from uncontrolled NPCST composition differences between study cohorts, and taxa identified as disease-associated may instead reflect NPCST-driven compositional shifts. This NPCST-aware framework constitutes a methodological advance: it redefines the analytical baseline from which disease effects are measured, because conventional unstratified comparisons overlook the distinct ecological states that structure the nasopharyngeal microbiome.

To test this hypothesis, we conducted a comprehensive meta-analysis to date of the nasopharyngeal microbiome, encompassing 28 cohorts with 7,790 uniformly reprocessed 16S rRNA samples. Shotgun metagenomic approaches remain impractical at this body site because high host-content and low-biomass samples overwhelm microbial signal [38], making 16S rRNA gene sequencing the dominant and most tractable approach for nasopharyngeal microbiome research. We employed a stringent three-stage decontamination pipeline that systematically removed environmental and reagent-derived microbial families, ensuring that the retained microbial composition reflects genuine human-associated signal without compromising overall community structure (**Methods; Supplementary File**). With curated data, we identified six robust and reproducible NPCSTs, and using nested mixed-effects models, we show that NPCST identity, not disease status, drives the diversity patterns that have generated contradictory conclusions across the literature. We then applied NPCST-aware differential abundance testing via ANCOM-BC2 to SARS-CoV-2 as a case study, confirming 13 reproducible taxa that survive NPCST adjustment. SPIEC-EASI co-occurrence networks independently demonstrate that structurally central hub genera are preferentially depleted during infection, with persistent anaerobes serving as the conserved backbone across NPCSTs. To bridge nasopharyngeal microbiome research and clinical assessment, we developed and externally validated the Nasopharyngeal Microbiome Health Index (NMHI), a continuous community-level wellness metric that achieves high performance spanning diverse respiratory conditions. Finally, we further confirmed these findings using external validation cohorts alongside an independently generated discovery cohort.

Our results establish NPCSTs as a ubiquitous and common organizing principle of the nasopharyngeal microbiome, resolving previously contradictory findings while uncovering genuine disease-associated signals.

## Materials and methods

### Study screening and metadata evaluation

Our meta-analysis followed PRISMA guidelines (**Supplementary File** for checklist)[39]. We systematically searched PubMed [40] and Embase [41] on May 14, 2024, using the query: ("nasopharyngeal") AND ("microbiome" OR "microbiota"). Two independent reviewers applied exclusion criteria listed in Figure 1B. Initial screening of 710 publications yielded 227 for full-text review, of which 63 reported nasopharyngeal swab 16S rRNA gene sequencing data. After evaluating data downloadability, processing, and background decontamination, we retained 28 studies comprising 7,790 samples. This curation process was independently described by a companion study (Bustos et al.[42]), which documents the inconsistencies in nasopharyngeal microbiome reporting that necessitated excluding the majority of screened studies.

**Figure 1.**
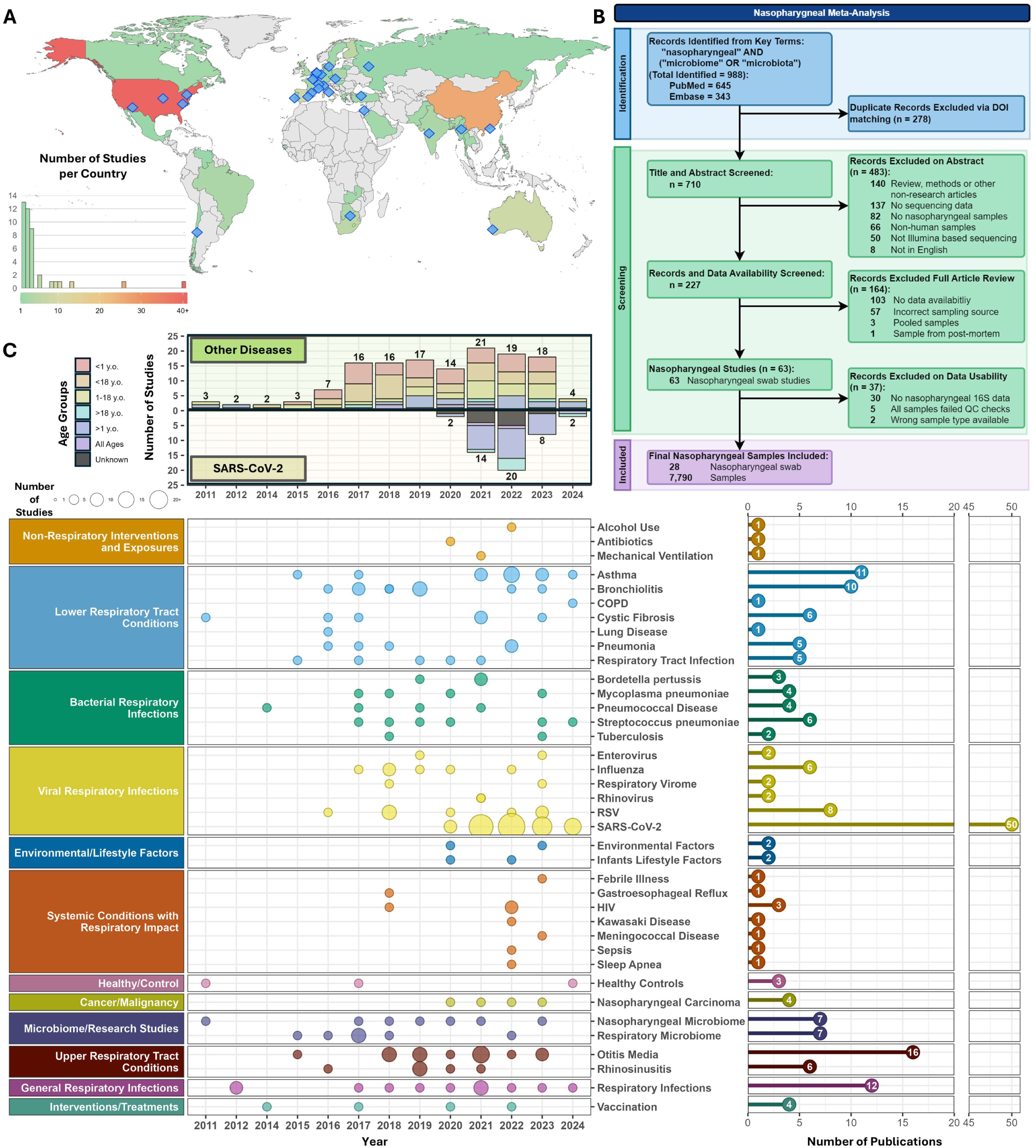
Global distribution and temporal trends of nasopharyngeal microbiome studies. A. World map displaying geographic distribution of publications examining human nasopharyngeal samples (n=181 across 42 countries). Color intensity indicates publication frequency per country (green to red gradient, 0-40+ studies). Blue diamonds mark locations of the 28 studies with downloadable datasets. B. PRISMA flow diagram for the systematic review. C. Temporal distribution of nasopharyngeal microbiome research by topic and age category (2011-2024). Studies are categorized into 10 high-level research topics encompassing 39 specific conditions, with bubble size proportional to annual study count and color-coded by research topic. Studies addressing multiple topics are counted in each relevant category. Upper panel: stacked bar plot depicting distribution of study publication years with colors distinguishing studies by age cohort studied. Studies are separately counted by condition (bottom: SARS-CoV-2 infection, top: other conditions). Right panel: total number of publications per condition.

### 16S data processing

Raw 16S rRNA gene sequencing datasets (**Table 1**) were processed using a standardized pipeline: automated hypervariable region identification via an in-house kmer hashing protocol built upon the minHaSH algorithm [43] and primer detection, then quality filtering and denoising with DADA2 [44] v1.22.0 and taxonomic classification with SILVA v138.2 [45] (80% bootstrap). Amplicon Sequence Variants (ASVs) were aggregated at the lowest assigned rank with samples required ≥ 5,000 family-level reads post-decontamination, and studies required ≥ 50% sample retention (**Supplementary File**). Phylogenetic reconstruction methods are detailed in the **Supplementary File**.

**Table 1.** Meta-analysis study characteristics. Each row shows the details of one of the public studies, including the disease/264 condition of interest, reference, population age range, PMCID, data source (NCBI BioProject Identifier), total samples available for download, hypervariable region (HVR), samples analyzed and passing quality control (QC), and cohort format.

| Disease/Condition of Interest | Reference | Age Range (Years)<sup>†</sup> | PMID | NCBI BioProject | Samples Available for Download | Hypervariable Region | Samples Passing QC | Samples Passing QC [%] | Cohort Type |
| --- | --- | --- | --- | --- | --- | --- | --- | --- | --- |
| Respiratory viruses | Edouard et al., 2018 [60] | 0-94 | 30033505 | PRJEB14780 | 451 | V3-V4 | 307 | 68.07% | Main |
| Chronic Rhinosinusitis | De Boeck et al., 2019 [34] | 18-65 | 31776238 | PRJEB30316 | 172 | V4 | 151 | 87.79% | Main |
| Otitis Media | Jörissen et al., 2021[61] | ~0-18 | 33879499 | PRJEB33591 | 114 | V4 | 113 | 99.12% | Main |
| Cystic Fibrosis | Pittet et al., 2021 [62] | <18 | 33247928 | PRJEB41059 | 5 | V3-V4 | 5 | 100.00% | Main |
| Pneumococcal Colonization on HIV exposed/unexposed | Kelly et al., 2022 [33] | <1 | 34511605 | PRJEB41741 | 156 | V3-V4 | 121 | 77.56% | Main |
| Influenza-like Illness | Shetty et al., 2022 [63] | ~63.5-77.6 | 35115596 | PRJEB46215 | 1,072 | V4 | 878 | 81.90% | Main |
| Asthma | Thorsen et al., 2023 [64] | <1 | 37863895 | PRJEB58215 | 290 | V3-V4 | 268 | 92.41% | Main |
| Meningococccemia | Bozan et al., 2023 [65] | <3 | 37370879 | PRJEB59932 | 49 | V3-V4 | 25 | 51.02% | Main |
| Tuberculosis | Ruiz-Tagle et al., 2023 [66] | ~32-39 | 37147354 | PRJEB61078 | 82 | V1-V3 | 24 | 29.27% | Main |
| Otitis Media | Lappan et al., 2018 [26] | 0.108 - 0.267 | 29458340 | PRJNA388707 | 195 | V3-V4 | 181 | 92.82% | Main |
| HIV and Tuberculosis | Htun et al., 2018 [67] | Not available | 30364728 | PRJNA432583 | 33 | V3-V4 | 31 | 93.94% | Main |
| Enterovirus D68 | Bowers et al., 2019 [68] | 1.5-17 | 30670612 | PRJNA474932 | 6 | V3-V4 | 4 | 66.67% | Main |
| SARS-CoV-2 | Ventero et al., 2021 [69] | 13-94 | 33815323 | PRJNA673585 | 74 | V3-V4 | 74 | 100.00% | Main |
| SARS-CoV-2 | Merenstein et al., 2021 [36] | 36-94 | 34399607 | PRJNA683617 | 267 | V1-V2 | 245 | 91.76% | Main |
| SARS-CoV-2 | Braun et al., 2021 [70] | 30-68 | 33903709 | PRJNA688646 | 55 | V4 | 30 | 54.55% | Main |
| SARS-CoV-2 | Hurst et al., 2022 [16] | 4-17 | 35247047 | PRJNA703574 | 285 | V4 | 260 | 91.23% | Main |
| Exacerbated Wheezers and Asthmatics | Aydin et al., 2021 [71] | 0.25-17 | 35386977 | PRJNA714100 | 64 | V3-V4 | 64 | 100.00% | Main |
| Respiratory Tract Infection | Piters et al., 2022 [31] | 0-1 | 35058634 | PRJNA740120 | 1,156 | V4 | 1,142 | 98.79% | Main |
| SARS-CoV-2 | Galeeva et al., 2022 [72] | Not available | 34977286 | PRJNA751478 | 335 | V3-V4 | 229 | 68.36% | Main |
| SARS-CoV-2 | Ventero et al., 2022 [73] | 52-80 | 34963638 | PRJNA754005 | 177 | V3-V4 | 171 | 96.61% | Main |
| SARS-CoV-2 | Crovetto et al., 2022 [9] | 27.3-35.8 | 35927569 | PRJNA777915 | 76 | V3-V4 | 75 | 98.68% | Main |
| Environmental Impact | Karampatsas et al., 2023 [74] | 20-69 | 37525095 | PRJNA788869 | 44 | V4 | 41 | 93.18% | Main |
| SARS-CoV-2 | Ferrari et al., 2022 [75] | ~33-57 | 35873175 | PRJNA839581 | 54 | V3-V4 | 51 | 94.44% | Main |
| Lower Respiratory Tract Infection | Koenen et al., 2023 [76] | 2-6 | 37950996 | PRJNA897835 | 358 | V4 | 265 | 74.02% | Main |
| SARS-CoV-2 | Kim et al., 2023 [12] | 37-77 | 36583481 | PRJNA912320 | 66 | V1-V2 | 58 | 87.88% | Main |
| Pneumococci Carrier | Paulo et al., 2023 [32] | 25-50 | 37658443 | PRJNA962709 | 177 | V4 | 152 | 85.88% | Main |
| SARS-CoV-2 | Romani et al., 2024 [13] | <17.7 | 38289047 | PRJNA993438 | 106 | V3-V4 | 99 | 93.40% | Main |
| Healthy | Odendaal et al., 2024 [29] | 0-87 | 39094567 | PRJNA997934 | 2,799 | V4 | 2,726 | 97.39% | Main |
| SARS-CoV-2 | Galeana-Cadena et al., 2024 [10] | 20-87 | 38826746 | PRJNA981220 | 40 | V4 | 40 | 100.00% | External |
| RSV | Kristensen et al., 2024 [30] | < 1 | 39642873 | PRJNA914884 | 1,537 | V4 | 1,533 | 99.74% | External |
| †The symbol "~" indicates that exact ranges were not provided in the original source; values represent approximations based on reported means ± standard deviations or other available metrics. |  |  |  |  |  |  |  |  |  |

### Nasopharyngeal background decontamination protocol

Because negative controls were unavailable for most studies, we developed a three-stage decontamination pipeline integrating LLM-assisted screening (Claude [46] implementing triplicate custom queries per family, with disagreements among the triplicate queries resolved by expert override), expert curation, and statistical validation (**Figure 2A** and **Supplementary File**). Assessment was performed at the family level, the complete list of flagged environmental and reagent-derived contaminant families is provided in **Table S3**. To verify that filtering did not distort true community relationships, pre- and post-decontamination ordinations were compared using Procrustes and Mantel correlation tests (**Supplementary File**).

**Figure 2.**
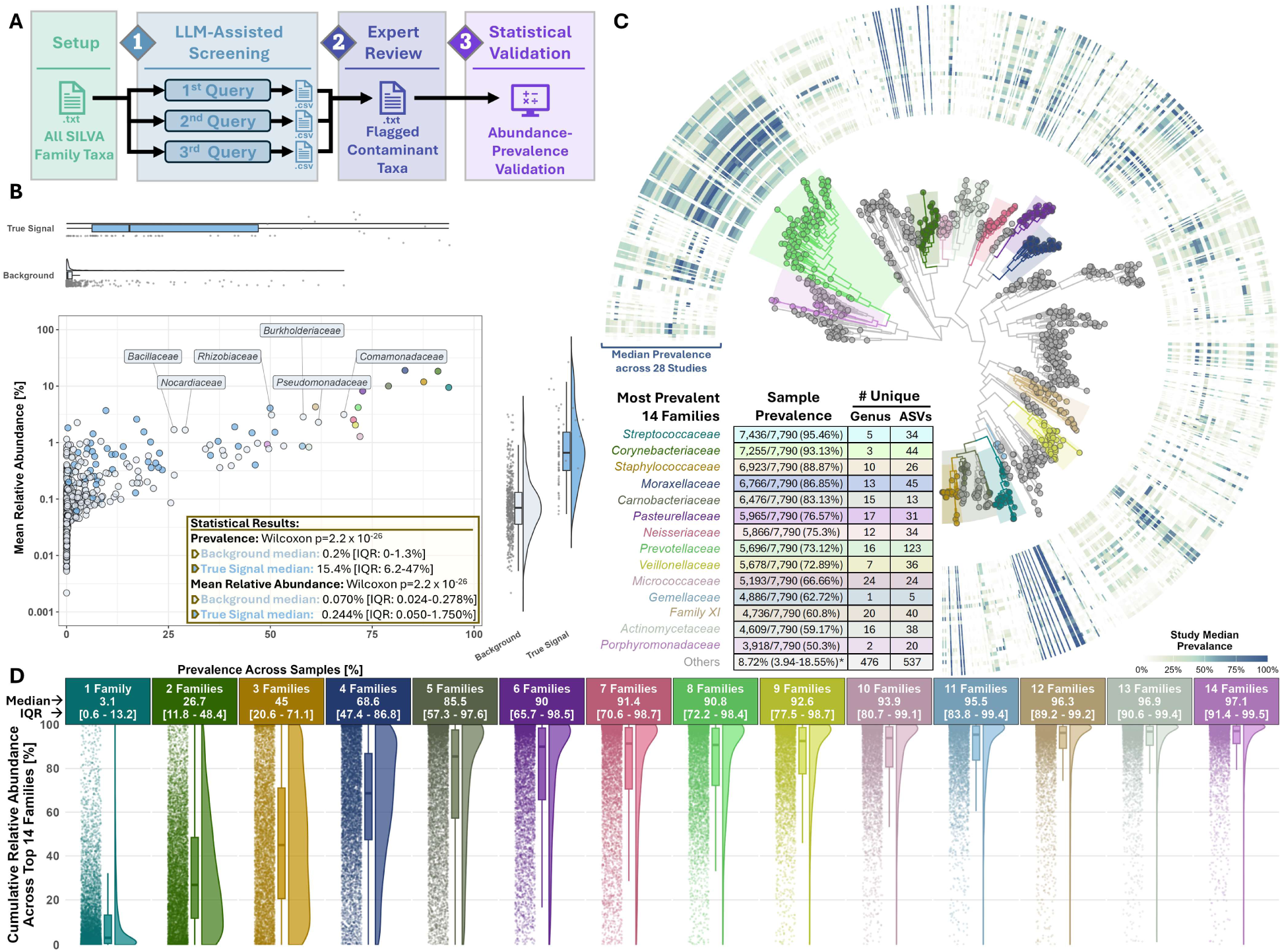
Three-stage contamination assessment and family-level abundance patterns in decontaminated nasopharyngeal microbiome datasets. **A.** Three-stage contamination identification workflow: (1) LLM-assisted screening of the 596 SILVA-annotated bacterial families detected, (2) expert review to yield initial taxon contamination classifications, and (3) statistical validation via abundance-prevalence analysis, establishing a nasopharyngeal-specific contaminant reference database. **B**. Scatter plot depicting the sample prevalence [%] (x-axis) and median relative abundance [%] (y-axis) of bacterial families colored by contamination status. Axis marginal distributions for contaminants (white) and true signals (i.e., authentic nasopharyngeal families; blue) are shown as boxplots with adjacent jittered data points with Wilcoxon rank-sum statistical analysis results shown in the scatter plot. Selected background families with atypically high prevalence (≥25%) and abundance (≥1%) are labeled. The top 14 prevalent bacterial families (21.9% of the 64 authentic nasopharyngeal families) classified as true signals are separately colored and further described in panels C-D. **C**. Maximum likelihood phylogenetic tree of 1,091 amplicon sequence variants (ASVs) from the representative V3-V4 hypervariable region (**Methods**), with the top 14 prevalent families highlighted by colored rectangular backgrounds organized by phylogenetic clades; overlapping colors denote shared ancestry. The outer heatmap displays the prevalence of the most-refined taxonomic classification associated with each ASV across the meta-analysis dataset (28 independent studies). **D**. Sample cumulative relative abundance distributions of the top 14 prevalent families, beginning with the most prevalent family (left) and with each subsequent panel showing the accumulation upon adding the next most prevalent family. Each panel displays the family abundance distribution summary statistics (IQR and median) and left-to-right as a jittered data points, boxplot, and density curve.

### Nasopharyngeal community state types (NPCST) classification

A minimum relative abundance threshold of 0.01% was applied at the most refined taxonomic resolution before aggregation to genus level. Taxa resolved only to family level were retained with the prefix "f_". Nasopharyngeal microbiome profiles were hierarchically clustered using Ward’s minimum variance method as established by France et al. [47]. Clustering was performed on pairwise Bray-Curtis dissimilarities computed using vegan v2.6-8 [48]. Optimal cluster number was determined by silhouette score analysis (cluster v2.1.6) [49], and confirmed through leave-one-study-out (LOSO) cross-validation, which systematically removed each study and regenerated clusters from the remaining 27. Cluster membership consistency was evaluated using the Adjusted Rand Index (ARI), Jaccard similarity, and silhouette scores across k = 2-20. K = 6 was selected as the value achieving the highest median performance across all three metrics (**Figure 3A** and **S5**). Two machine learning classifiers (Random Forest and SVM) were trained to predict NPCST assignment from genus-level profiles, providing a reproducible framework for future studies to classify nasopharyngeal samples (**Supplementary File** and **Figure S4**). Following NPCST establishment, PERMANOVA was performed using vegan adonis2 to test community composition differences among NPCSTs, with age, hypervariable region, sex, and study as covariates. Moreover, the Unsupervised Learning based Definition of the Rare Biosphere (ulrb) [50] method was employed to define NPCST-specific taxa prevalence categories (abundant, intermediate, rare), using data-driven abundance cutoffs, providing the taxonomic framework for the heatmap visualization (**Figure 3D**) and subsequent key taxa selection.

**Figure 3.**
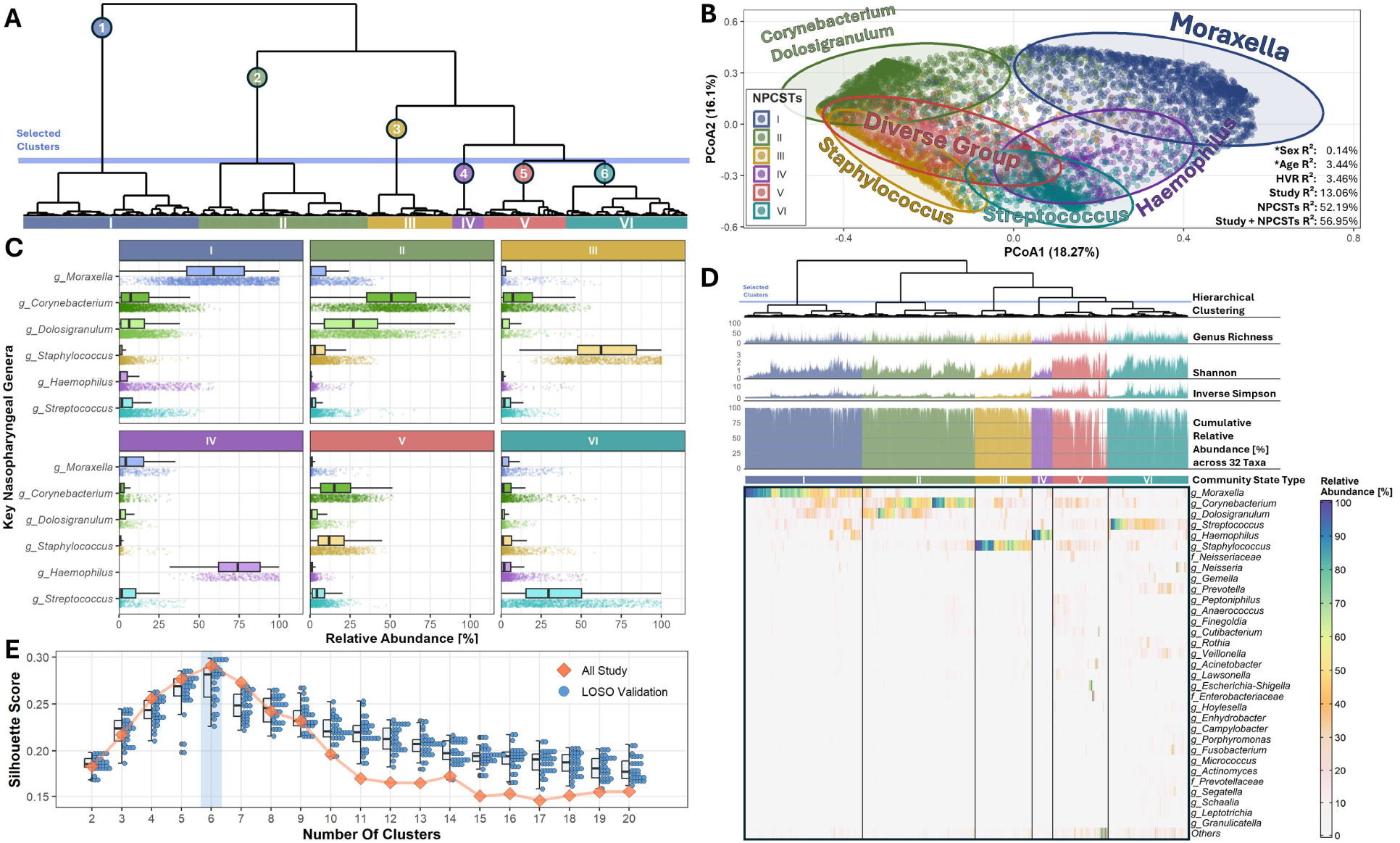
Nasopharyngeal Microbiome Community State Type (NPCST) Analysis. **A.** Hierarchical clustering of nasopharyngeal microbiomes via Ward’s Method using pairwise Bray-Curtis dissimilarities. The dendrogram was cut at the blue horizontal line to yield six distinct NPCSTs colored and described further in panels B-E (I=blue, II=green, III=yellow, IV=purple, V=red, VI=teal). **B.** Principal coordinates analysis (PCoA) based on pairwise Bray-Curtis dissimilarities with each NPCST indicated by a colored ellipse representing 95% confidence intervals. The PERMANOVA R² of key sample parameters (sex, age, hypervariable region (HVR), study, NPCST) were individually calculated with a combined mode (Study + NPCSTs) displayed. **C.** Relative abundance [%] distributions of key genera (*g_Moraxella, g_Corynebacterium, g_Dolosigranulum, g_Staphylococcus, g_Streptococcus,* and *g_Haemophilus*) across the six NPCSTs represented by vertically adjacent boxplots and jittered sample data points. **D.** Heatmap displaying relative abundances of 32 ulrb-identified taxa with prevalence ≥50% in at least one NPCST (**Figure S6-11**). Above the heatmap top-to-bottom are the sample hierarchical clustering dendrogram, genus richness (i.e., number of genera detected), alpha diversity (Shannon diversity and Inverse Simpson diversity), and cumulative relative abundance of the 32 ulrb-identified taxa. For visualization purposes, Inverse Simpson diversity is capped at 10, genus richness at 100, and Shannon diversity at 3. **E.** Silhouette score validation for determining the optimal number of clusters (x-axis), where the leave-one-study-out validation silhouette score distributions are shown as boxplots adjacent to their respective jittered data points (y-axis). The connected orange data points represent silhouette scores calculated using the complete meta-analysis dataset (all 28 independent studies).

### Diversity paradox analysis

Unless otherwise noted, all analyses below use only cross-sectional samples (one sample per subject) to avoid bias from repeated measures. To test whether disease or NPCST architecture drives alpha diversity patterns, we fitted three nested linear mixed-effects models [51] to Shannon diversity across 5,464 cross-sectional samples with clear disease and control status and NPCST assignment (ambiguous cross-sectional categories, e.g., 1-month-old infants, were excluded). Model C additionally required complete age and sex data, restricting that model to 3,989 of these samples (73%):

- Model A: Shannon ∼ DiseaseStatus + (1|BioProject)
- Model B: Shannon ∼ DiseaseStatus + NPCST + (1|BioProject)
- Model C: Shannon ∼ DiseaseStatus + NPCST + Age + Sex + (1|BioProject)

Disease status was coded as a binary variable (control vs. disease). Significance of fixed effects was assessed using lmerTest [52] and effect attenuation was quantified as the percentage reduction in the disease coefficient between Models A and B. NPCST-disease enrichment was quantified using multinomial logistic regression (mclogit v0.9.6 [53] ) with NPCST membership as outcome and disease category (bacterial infection, viral infection, control) as predictor, with BioProject included as a random effect to account for study-level heterogeneity. Odds ratios with 95% confidence intervals were derived from marginal probability contrasts (emmeans v1.10.5 [54]).

### ANCOM-BC2 differential abundance testing with LOSO cross-validation

To distinguish genuine disease-associated taxa from NPCST-driven compositional shifts, we applied ANCOM-BC2 [55] to SARS-CoV-2 samples using two complementary models: (1) with BioProject as covariate (unadjusted for NPCST), and (2) with BioProject and NPCST as covariates (NPCST-adjusted). Coefficient attenuation between models quantified the proportion of signal attributable to NPCST composition versus genuine disease association. Both models used Benjamini-Hochberg (BH) correction (adjusted p < 0.05) and a two-tier prevalence filter: global prevalence ≥5% AND per-group prevalence ≥10% within comparison groups. For the within-NPCST diversity comparison (**Figure 5C**), this analysis used a cross-sectional-only SARS-CoV-2 cohort of 7 studies (918 samples: 261 controls, 657 cases).

To assess taxon reproducibility independent of any single cohort, we performed leave-one-study-out (LOSO) cross-validation across an expanded cohort of 10 SARS-CoV-2-contributing studies (1,100 samples; 812 cases, 288 controls). This expanded cohort additionally included baseline timepoints from longitudinal studies and an external validation cohort (PRJNA981220) not included in the cross-sectional cohort. For each iteration, the entire held-out study was removed and ANCOM-BC2 was re-run on the remaining samples with identical parameterization. Reproducible taxa were defined as those with |mean LOSO fold change| ≥0.4 and directional consistency ≥90% (i.e., same direction in ≥9 of 10 LOSO iterations).

### Co-occurrence network analysis

SPIEC-EASI (v1.1.2)[56] was employed to infer microbial associations globally and within individual NPCSTs, using the neighborhood selection ("mb") algorithm on genus-aggregated abundance matrices. Only cross-sectional samples were analyzed to prevent bias from repeated measures. Taxa required ≥ 1% prevalence and >10 reads. Hub taxa were identified through k-core decomposition (igraph v2.1.1 [57]), with the upper quintile (top 20%) defining structurally important network components. To test whether network topology predicts disease sensitivity, we compared absolute ANCOM-BC2 log-fold changes between hub and non-hub genera using Wilcoxon rank-sum tests, stratified by oxygen requirement (aerobic vs. anaerobic).

### Nasopharyngeal microbiome health index development

Building upon the Gut Microbiome Wellness Index 2 (GMWI2) framework [58] as a conceptual foundation, we developed the NMHI through LASSO-penalized logistic regression with balanced class weights and binary presence/absence encoding at 0.01% relative abundance threshold, applied only to cross-sectional samples. Beyond GMWI2 conceptual foundation, the NMHI introduces three NP-specific advances: Leave-One-NPCST-Out (LONO) cross-validation for lambda selection, NPCST-stratified performance thresholds, and NP-specific feature selection. The Leave-One-NPCST-Out (LONO) cross-validation evaluated a range of lambda (0.0001-0.03), followed by 10-fold CV across five prevalence thresholds between 0% and 20%.

The five-stage methodology comprised: (1) LONO CV for lambda optimization; (2) performance evaluation across prevalence thresholds, selecting 5% for optimal performance-interpretability balance; (3) final model training to extract coefficients, with NMHI = intercept + sum of present taxa coefficients; (4) NMHI healthy/disease boundary optimization (-5 to 5 with 0.1 increments) maximizing balanced accuracy globally and per-NPCST; (5) external validation on 699 independent samples spanning healthy controls, SARS-CoV-2 severity levels, lower respiratory tract infections, and critical illness. Complete methodology is provided in the **Supplementary File**.

### External Validation dataset

Two additional BioProjects released after the initial extraction cutoff yielded 1,573 samples for NPCST classifier validation. NMHI external validation used an independent set of 699 samples. BioProject accessions and processing details are provided in **Table 1** and **Supplementary File**.

### Discovery cohort and PCR panels

To validate the full analytical framework in an independent clinical setting, we processed 147 remnant respiratory panel specimens via 16S rRNA gene sequencing, using the same V3-V4 region amplified on an Illumina MiSeq (Illumina, San Diego, CA, USA) and processed through the identical DADA2 and SILVA v138.2 pipeline as the meta-analysis. As these samples were derived from clinical diagnostic remnants, matched healthy controls were not available. A separately drawn, partially overlapping set of 124 of these specimens additionally underwent multiplex RT-PCR (Seegene® Novaplex™ panels). NPCST classification was performed using the pre-trained Random Forest model, and NMHI scores were calculated by applying the 5% prevalence model coefficients (197 features) to binary presence/absence matrices constructed from the discovery data. Reference pathogen detection was established through total RNA-sequencing with Kraken2-based taxonomic classification [59] for comprehensive respiratory pathogen detection on these 147 samples.

### Statistical methods

Full statistical procedures, software, and effect-size reporting conventions are detailed in the **Supplementary File**.

## Results

### A Standardized Nasopharyngeal Microbiome Compendium

Following PRISMA guidelines (**Supplementary File**), we systematically screened 710 publications and identified 28 studies with downloadable 16S rRNA gene sequencing data, encompassing 7,790 samples after uniform bioinformatic reprocessing (**Table 1**; **Figure 1A-B**). Overall, nasopharyngeal microbiome research spanned 42 countries, with research accelerating after 2017 (**Figure 1C**). Post-2020 adult cohorts are almost exclusively SARS-CoV-2 studies, while non-SARS-CoV-2 research remains predominantly pediatric (<18 y.o.). Our final compendium covers diverse respiratory infections of bacterial and viral origin alongside healthy samples (**Table 1)**.

Because negative control samples were unavailable for most studies, we developed a three-stage decontamination pipeline integrating LLM-assisted screening, expert curation, and statistical validation (**Methods, Supplementary File** and **Figure 2A**). This pipeline identified 532 of 596 detected bacterial families (89.3%) as environmental or reagent-derived contaminants, which exhibited significantly lower prevalence (median 0.2% vs 15.4%; p < 2.2 x 10-26) and abundance than authentic signal (**Figure 2B**). The resulting dataset retained 64 curated bacterial families, of which 14 core families colonized >50% of samples and comprised a median 97.1% of total abundance across all 7,790 samples, establishing the fundamental taxonomic architecture of the nasopharyngeal microbiome (**Figure 2C-D**).

### Characterization of nasopharyngeal NPCSTs

We performed NPCST analysis via hierarchical clustering of the 7,790 samples resulting in six distinct and stable NPCSTs (**Figures 3A**). These NPCSTs displayed characteristic ecological signatures: NPCST I (*Moraxella*-dominated), NPCST II (*Corynebacterium*-*Dolosigranulum* co-dominated), NPCST III (*Staphylococcus*-dominated), NPCST IV (*Haemophilus*-dominated), NPCST V (diverse, no dominant genus), and NPCST VI (*Streptococcus*-dominated) (**Figures 3B-C**). These NPCST keystone taxa were among the 14 core families (**Figure 2C-D**), and they were highly prevalent (detected in >80% of samples) and abundant within their respective NPCSTs (**Figures 2B**, **3C** and **S4**). More importantly, NPCSTs explained 52.19% of total variance from PERMANOVA in community composition, ∼4x higher than study-specific batch effects (13.06%) and far exceeding other covariates: HVRs (3.46%), age (3.44%), and sex (0.14%) (**Figure 3B**). This dominant biological signal shows that these six NPCSTs represent dominant ecological states in the nasopharyngeal environment.

These six NPCSTs demonstrated robust reproducibility across multiple validation approaches (**Supplementary File**) and exhibited community specific taxon abundance patterns (**Figure 3D**). Cluster number optimization using silhouette score, Adjusted Rand Index, and Jaccard similarity identified six clusters as optimal, with LOSO validation confirming k=6 as the point of maximum stability across all three metrics over the tested k=2-20 range (**Figures 3E** and **S5**). Furthermore, ulrb analysis independently validated dominant and sub-dominant taxa patterns, revealing 2-6 abundant taxa per NPCST with consistent detection in >80% of samples within each NPCST (**Figures S6-S11** and **Table S4**). Diversity metrics clearly differentiated the NPCSTs: NPCSTs I-IV represented lower-diversity range dominated by 1-2 genera, whereas NPCSTs V-VI comprised more diverse compositions with higher genus richness and diversity indices (**Figure 3D**).

We developed and validated machine learning models that enable standardized NPCST classification (**Supplementary File**), achieving high median accuracy using both Random Forest (96.6%) and Support Vector Machines (97.2%). The Random Forest NPCST classifier demonstrated robust external validation on 1,573 independent samples, achieving NPCST classifier AUCs >0.91 for NPCSTs I-IV and VI, though NPCST V showed lower performance (0.853), consistent with its compositional heterogeneity (**Figure S12**). These models show that failing to account for NPCST composition risks confounding future nasopharyngeal 16S microbiome studies. We further validated this classification framework using our own discovery set.

### Resolving the diversity paradox through NPCST stratification

To test whether conflicting diversity reports arise from uncontrolled NPCST composition, we fitted three nested linear mixed-effects models to Shannon diversity across cross-sectional samples with study as a random effect (**Figure 4A-B**). Within individual NPCSTs, pairwise Wilcoxon tests revealed that most disease-control comparisons were non-significant after BH correction with the few exceptions (NPCSTs II, IV, V for viral infection) showed small effect sizes relative to inter-NPCST differences (**Figure 4A**). Model A (disease only) explained negligible variance (marginal R² = 0.029), whereas Model B (adding NPCST) increased explanatory power 13-fold (R² = 0.382; ΔAIC = -1860) and attenuated disease coefficients by 92.5% (bacterial) and 93.5% (viral), rendering both fully non-significant (**Figure 4B**). Model C (adding NPCST, age and sex) provided no further improvement (R² = 0.385). Moreover, multinomial logistic regression with study stratification quantified NPCST-disease enrichment (**Figure 4C**). NPCST VI showed significantly elevated bacterial infection representation (p = 0.012) and reduced viral representation (p < 0.001), while NPCSTs III and IV showed the opposite pattern with viral infection enrichment (p = 0.021 and p = 0.028, respectively). These differential enrichment patterns explain why studies sampling populations with different NPCST compositions report contradictory diversity findings and establish that all comparative analyses of the nasopharyngeal microbiome require NPCST stratification or adjustment.

**Figure 4.**
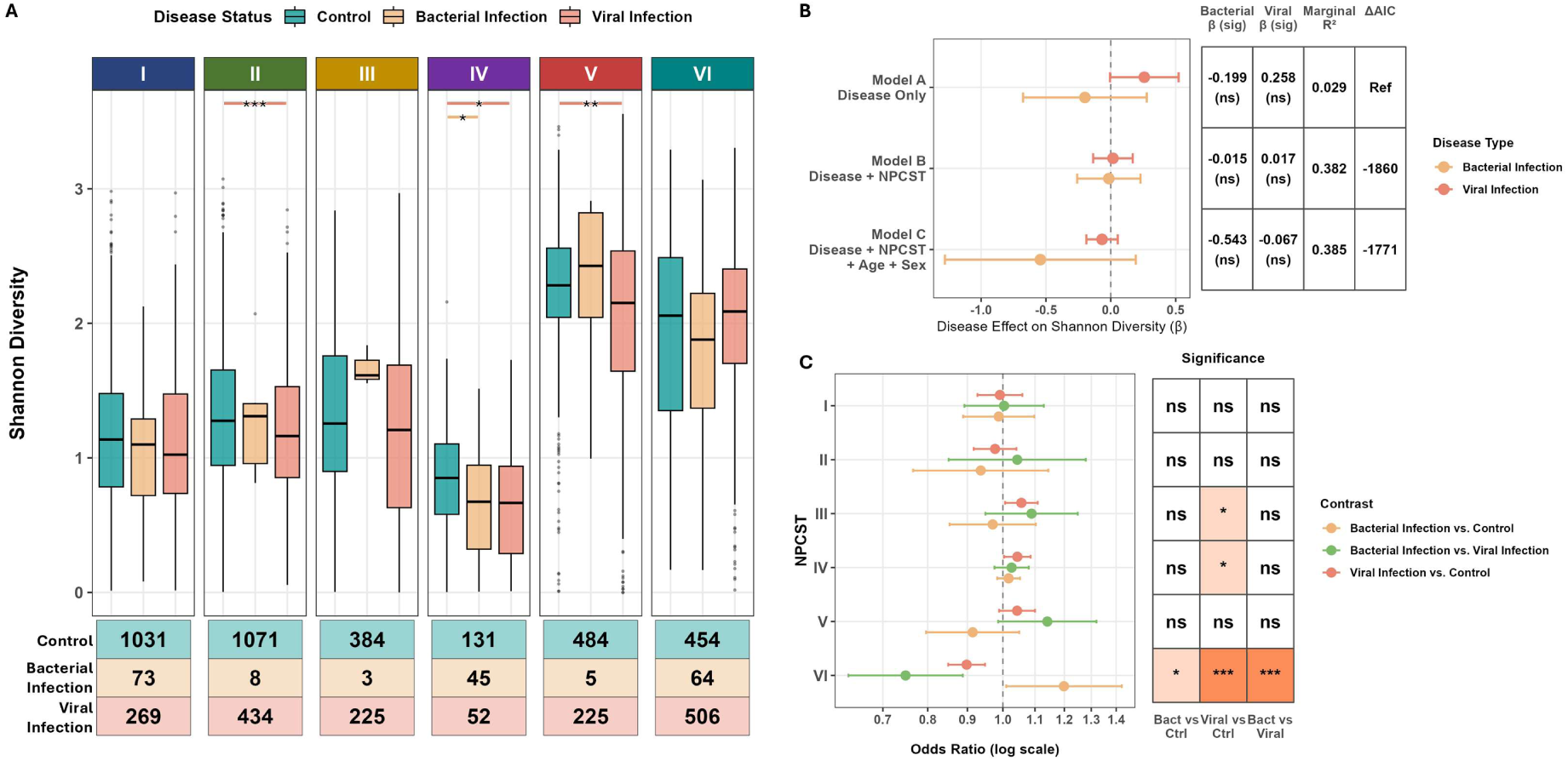
NPCST architecture resolves the diversity paradox and reveals disease-specific enrichment patterns. **A.** Shannon diversity across six NPCSTs stratified by disease status (Control, Bacterial Infection, Viral Infection) in 5,464 cross-sectional samples. Pairwise Wilcoxon rank-sum tests (BH-adjusted) are shown for significant comparisons. Sample sizes per group are displayed below. **B.** Disease effect attenuation across three nested linear mixed-effects models. Left: coefficient estimates (beta, 95% CI) for bacterial and viral infection effects on Shannon diversity. Right: model summary statistics including marginal R², ΔAIC, and significance. **C.** NPCST enrichment by disease type from multinomial logistic regression with study stratification. Left: odds ratios (95% CI, log scale) for each contrast. Right: significance tile (BH-adjusted; *p < 0.05, **p < 0.01, ***p < 0.001, ns = not significant).

**Figure 5.**
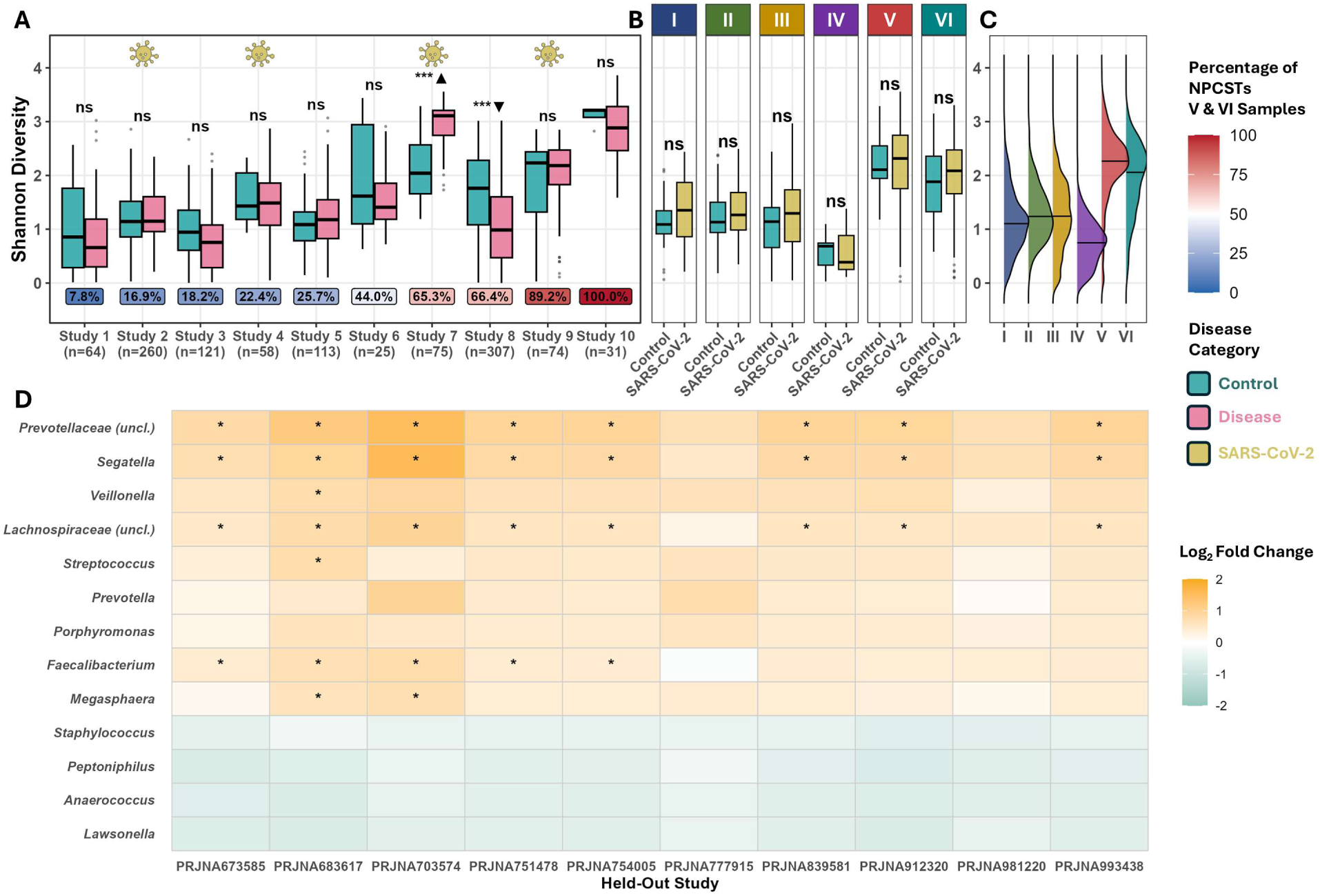
SARS-CoV-2 confirms NPCST-dependent diversity and reveals genuine disease markers. **A.** Study-level Shannon diversity comparisons between healthy controls and disease across all studies, ordered by proportion of high-diversity NPCSTs (V/VI). Four SARS-CoV-2-contributing studies are highlighted with virus symbols that are used for Panel B and C along with 3 additional SARS-CoV-2 only studies (n: 918 samples with 657 SARS-CoV-2 samples). **B.** Distributions of Shannon diversity by NPCST. **C.** Within-NPCST Shannon diversity comparisons between controls and SARS-CoV-2 patients pooled across 7 SARS-CoV-2 related studies (918 samples). **D.** ANCOM-BC2 LOSO CV heatmap showing log fold changes for 13 reproducible markers (|LFC| ≥ 0.4) across 10 held-out SARS-CoV-2 cohorts. BH-adjusted annotation: *p < 0.05, **p < 0.01, ***p < 0.001 and ns = not significant.

### SARS-CoV-2 confirms NPCST-dependent diversity and reveals genuine disease markers

Across all diseases from 10 studies, study-level diversity comparisons (healthy vs disease) showed heterogeneous patterns: one study reported significantly increased diversity, one reported decreased diversity, while the majority remained non-significant (**Figure 5A**).

These conflicting directions tracked directly with NPCST V/VI composition (**Figure 5A-B**), because samples can be dominated by high- or low-diversity NPCSTs, integrative analyses must account for this study-level compositional bias. When we refined the analysis to a disease-specific cohort using SARS-CoV-2, within-NPCST comparisons between controls and SARS-CoV-2 patients yielded no significant differences across any of the six NPCSTs after BH adjustment (**Figure 5C**). This is consistent with the apparent diversity heterogeneity observed at the study level arising predominantly from differential NPCST composition rather than genuine disease effects on alpha diversity.

To identify genuine disease-associated taxa independent of NPCST architecture, we applied ANCOM-BC2 with LOSO cross-validation across an expanded cohort of 10 SARS-CoV-2-focused studies, which additionally drew on baseline timepoints from longitudinal studies and an external validation cohort not included in the **Figure 5C** cross-sectional cohort above. This analysis identified 13 reproducible taxa with log fold change ≥0.4 and directional consistency ≥90%, revealing a bidirectional pattern. Obligate anaerobes typically reported in respiratory and gut communities (*Faecalibacterium*, unclassified *Lachnospiraceae*, unclassified *Prevotellaceae*, *Segatella*) [65, 77, 78] were consistently enriched in SARS-CoV-2 patients, while NP-resident hub genera (*Anaerococcus, Peptoniphilus, Lawsonella, Staphylococcus*) were depleted with 100% directional consistency across iterations (**Figure 5D**; see Co-occurrence networks section below). NPCST adjustment attenuated these signals by only 1.5%, confirming them as genuine disease markers independent of community state. Overall, these findings demonstrate that SARS-CoV-2 disrupts specific taxa without altering community-level diversity, explaining why unstratified analyses fail to detect genuine disease signals.

### Co-occurrence networks reveal structural backbone

To illustrate the core microbes within the nasopharyngeal space, we utilized SPIEC-EASI to construct NPCST-specific and global co-occurrence networks across cross-sectional samples (**Methods**). The results revealed sparse architectures in NPCSTs I-IV and dense, interconnected topologies in NPCSTs V-VI (**Figure 6A-F**). Anaerobic genera, particularly the *Peptoniphilus-Anaerococcus* hub pair, emerged as structural backbones across multiple NPCSTs, maintaining 23-81% prevalence despite the aerobic nasopharyngeal environment (**Figure 6G-H** and **S14**). Few hub genera were conserved across four or more NPCSTs, indicating a shared structural backbone that persists regardless of community architecture (**Figure 6H-I**).

**Figure 6.**
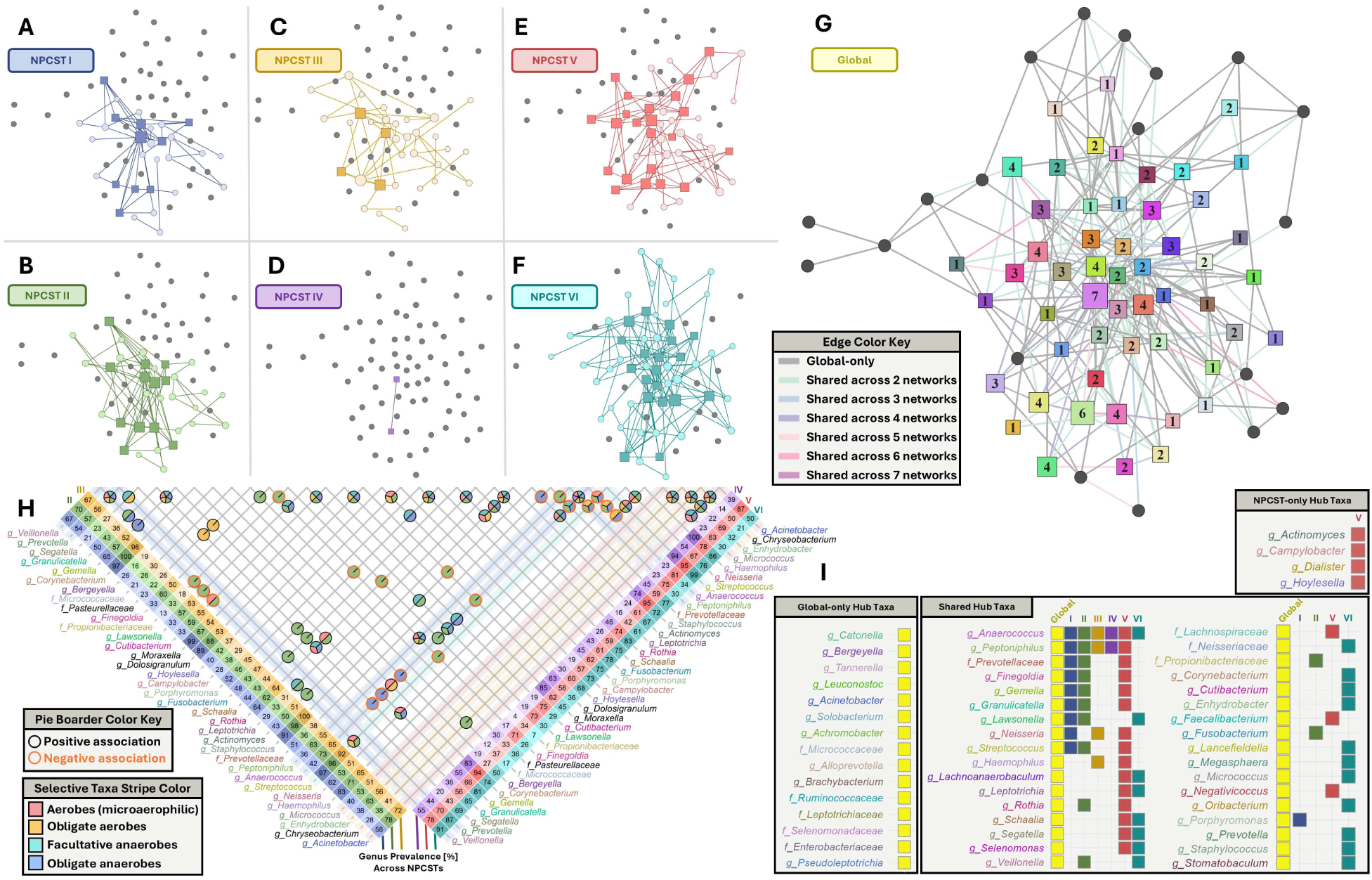
Taxon co-occurrence analysis of nasopharyngeal microbiome community state types (NPCSTs). A-F. NPCST-specific taxon co-occurrence networks derived from SPIEC-EASI analysis. Node colors correspond to NPCST-specific color schemes. Dark-colored squares represent k-core-determined hub taxa within each NPCST (**Methods**), while light-colored circles indicate non-hub taxa with associations. Gray nodes represent taxa present in other networks but lacking associations in the displayed NPCST. **G**. Global co-occurrence network integrating all samples across the six NPCSTs. Squares and circles denote hub taxa and non-hub taxa, respectively, with node size proportional to hub frequency across networks (labeled with number). Nodes (taxa) are also colored by hub category: global-only, NPCST-specific, or shared (global + NPCST). Edge colors indicate network sharing frequency, from gray (single network) to purple (all seven networks). All networks maintain consistent spatial positioning for direct comparison. **H**. Taxon-taxon associations for NPCSTs I–IV, with NPCSTs V–VI shown in **Figure S21C**. Pie charts represent pairwise genera connections, with segmentation indicating associations shared across multiple NPCSTs (color-coded by NPCST). Black and orange circles around the pie charts indicate positive and negative associations, respectively. Notably, all associations maintain consistent directionality across networks. Left and right axes display taxon prevalence percentages across NPCSTs. Selected taxa are highlighted with colored strips indicating oxygen requirements: aerobes (red), obligate aerobes (orange), facultative anaerobes (teal), and obligate anaerobes (blue). Taxon colors correspond to those shown in **panel I**. **I.** Legend displaying color-coded taxon names and their hub classification across network categories.

Network hub genera showed 2.71-fold larger absolute log-fold changes in SARS-CoV-2 compared to non-hub genera (Wilcoxon p = 0.008), with anaerobic hubs showing the strongest disruption (5.4-fold, p = 0.0007) (**Figure 6I**). Within NPCST VI, anaerobic hub disruption was bidirectional: NP-resident hubs were depleted while non-resident obligate anaerobes were enriched, a pattern masked in unstratified analyses (**Figure 5D**).

### Nasopharyngeal Microbiome Health Index bridges community ecology and clinical assessment

Besides the transient and stable microbes examined above, nasopharyngeal microbiomes are further complicated by poly-microbial colonization of obligatory and opportunistic microbes. More precisely, 4/6 NPCST keystone genera contain well-established respiratory pathogens (*Haemophilus influenzae*, *Streptococcus pneumoniae*, *Moraxella catarrhalis*, *Staphylococcus aureus*), yet these same genera define healthy community states. We verified poly-microbial presence via multiplex RT-PCR applied to 124 clinical NP specimens, which revealed that 66.9% harbored multiple pathogenic agents simultaneously (**Figure S15**). These observations complicate a binary disease classification approach that does not account for potential pathogens in the sample.

To counter this limitation and bridge the lack of clinical diagnostics for the nasopharyngeal microbiome, we developed the Nasopharyngeal Microbiome Health Index (NMHI) using LASSO-penalized logistic regression with binary presence/absence encoding on 5,435 cross-sectional samples (**Methods**). Leave-one-NPCST-out cross-validation achieved NMHI AUC = 0.886, with 10-fold CV confirming stable performance (NMHI AUC = 0.900; **Figure 7A**). The most explainable model with all-disease, all-taxa identified 98 disease-associated and 99 health-promoting taxa (**Figure 7B-C**). These findings demonstrate that nasopharyngeal wellness depends on the joint contribution of microbial presence across the community rather than abundance in any single organism, which avoids both the binary disease-classification problem above and any single NPCST’s dominance bias. The NMHI captures this contribution as a continuous health spectrum. Notably, seven of nine hub genera conserved across ≥4 NPCST networks were independently selected as NMHI features, with *Anaerococcus* species among the health-promoting signals. NMHI accounts for species-level duality within hub genera, e.g., *Haemophilus* at the genus level promotes health, yet *Haemophilus influenzae* specifically drives disease (**Figure 7D**). Across all six NPCSTs, NMHI distributions clearly separated healthy from diseased samples at a global threshold of 0.2 (Cohen’s d = 1.661-2.012; Wilcoxon p < 0.001), demonstrating that NMHI performance generalizes across community architecture (**Figure 7E**). External validation on 699 independent samples spanning healthy controls, SARS-CoV-2 severity levels, lower respiratory tract infections, and critical illness demonstrated discrimination across eight disease populations (median Cohen’s d = 2.179; **Figure 7F**), with an overall NMHI AUC of 0.922 and balanced accuracy of 81.5%, with NPCST-specific NMHI models performing comparably (NMHI AUC: 0.848-0.953; **Figure 7G**).

**Figure 7.**
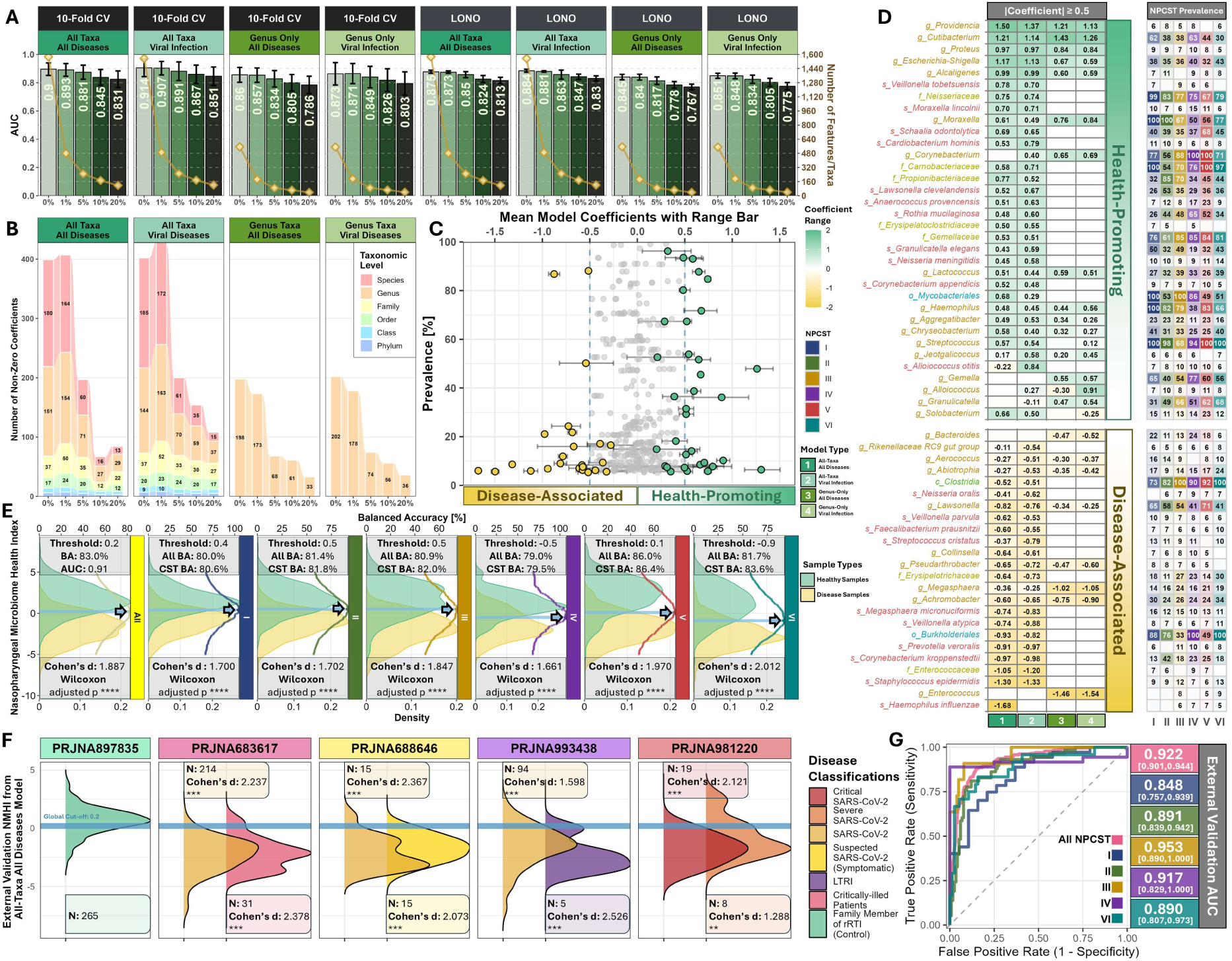
Nasopharyngeal Microbiome Health Index (NMHI) Validation. **A.** NMHI model performance evaluated using 10-fold cross-validation (CV) and Leave-One-NPCST-Out (LONO) validation across different microbial feature prevalence thresholds (0%, 1%, 5%, 10%, and 20%). Green bars show model median AUC values (left y-axis) with error bars indicating standard deviation across folds (CV) and NPCSTs (LONO). The connected yellow data points indicate the total features used for model training (right y-axis) at each prevalence threshold (x-axis). Each panel shows the results from the combination of validation type (CV or LONO) and the type of model generated, i.e., input microbial feature set used (all taxa or genus-level only) and model classification type (all conditions or viral infections)**. B.** Stacked bar plots showing the numbers of non-zero coefficients in final models trained on the complete dataset, stratified by prevalence threshold and model type. Features are colored by taxonomic resolution (species to phylum) with values ≥10 labeled on the bars. **C.** Mean coefficients (x-axis) and prevalence (y-axis) for each taxon across all four models with error bars indicating the minimum-maximum range. Taxa with an absolute coefficient ≥0.5 (vertical blue dashed lines) in at least one model are highlighted based on whether they are health-promoting (green; positive coefficients) or disease-associated (yellow; negative coefficients). Gray points represent taxa with absolute coefficients <0.5 across all models. **D.** Heatmaps showing the coefficient magnitudes for high-impact taxa (|coefficient| ≥0.5) across the four models (left) and corresponding prevalence distributions across NPCSTs (right), separately shown for health-promoting (top) and disease-associated (bottom) taxa. **E.** NMHI distributions of healthy controls (green) versus diseased samples (yellow) across all NPCSTs (left) and individual NPCSTs (left-to-right: I-VI). Horizontal dashed lines indicate optimal classification thresholds with arrows pointing to the corresponding maximum balanced accuracy values. Threshold values and balanced accuracy (BA) metrics are shown above each panel. Balanced accuracy is shown using the global threshold of 0.2 shown in the left-most panel ("All BA") and using the NPCST-specific threshold (“CST BA”). AUC is also shown above the left-most panel. Cohen’s d effect sizes and FDR-adjusted Wilcoxon p-values are shown below each panel. **F.** External validation of NMHI across five independent studies, comparing healthy controls with samples from patients with different conditions, including various SARS-CoV-2 infection severities, lower tract respiratory infection (LTRI), and critical illness patients. Cohen’s d values and FDR-adjusted Wilcoxon p-values are shown for each comparison with study BioProject accession shown above. **G.** External validation receiver operating characteristic (ROC) curves using global (All) and NPCST-specific thresholds, with AUCs and 95% confidence intervals displayed on the right. An ROC curve and AUC analysis was excluded for NPCST V since it did not contain any health controls. Asterisks denote statistical significance from Wilcoxon rank-sum tests (***p<0.001, **p<0.01).

### Discovery cohort confirms NPCST reproducibility and NMHI applicability

To validate the full framework in an independent clinical cohort, we processed 147 clinical nasopharyngeal swabs with known detection of RNA, DNA, and bacterial pathogens (**Methods** and **Supplementary File**). NPCSTs explained 52.3% of total variance (PERMANOVA p = 0.001) while data source contributed negligibly (R² = 0.09%; **Figure 8A**). Pre-trained Random Forest classifiers achieved 80.3% accuracy **(Figure 8B**) with misclassification concentrated in NPCST V (19/29 errors), consistent with this community state’s heterogeneous definition, as clinical specimens may harbor diverse profiles not captured in the published cohort training data. Excluding NPCST V, accuracy improved 91.5%. Within-NPCST Shannon diversity distributions overlapped substantially with meta-analysis baselines, confirming reproducible NPCST-specific diversity ranges across independent cohorts (**Figure 8C**). NMHI scores correlated with respiratory pathogen burden quantified by total RNA-Seq results (**Supplementary File**), with all bacterial and viral pathogen-positive samples showing median scores below the 0.2 global threshold from the NMHI model (**Figure 8D**). Consistent with its highest representation in the training data, SARS-CoV-2 samples showed the lowest median NMHI score. Overall, NMHI scores distributed consistently across all NPCSTs, confirming that NMHI performance generalizes across community architecture.

**Figure 8.**
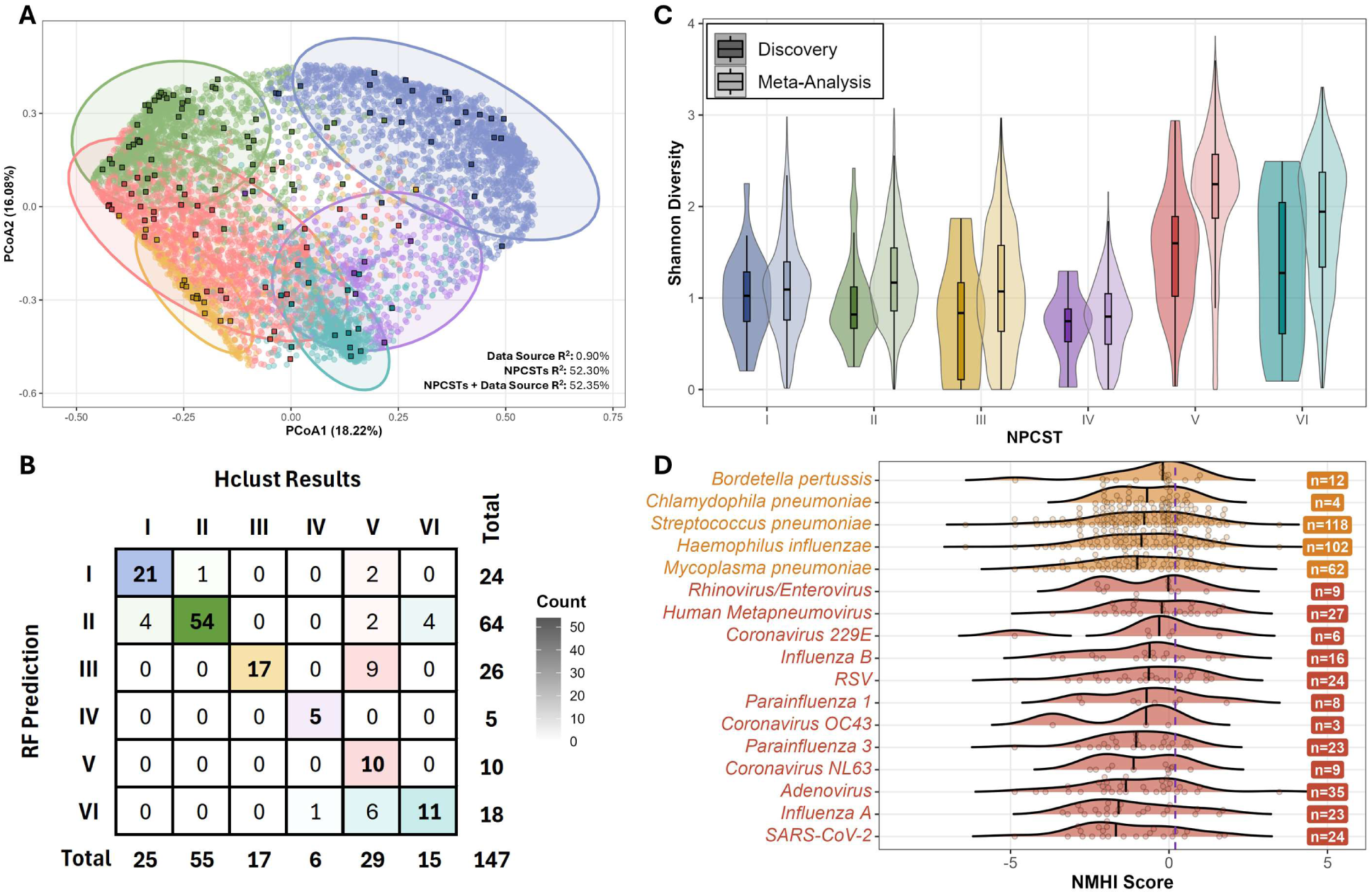
Discovery cohort validates NPCST framework and NMHI clinical utility. **A.** PCoA of Bray-Curtis dissimilarity showing 147 discovery specimens (squares) integrating within 7,790 meta-analysis samples (circles) by NPCST. PERMANOVA R-squared is displayed. **B.** Random Forest NPCST classification accuracy for the 147 discovery specimens using the pre-trained classifier comparing to the hierarchical clustering methods used to establish the 6 NPCSTs (**Methods**). **C.** Shannon diversity distributions per NPCST for discovery (darker color) and meta-analysis (lighter color) cohorts, confirming reproducible NPCST-specific diversity ranges. **D.** NMHI score distributions by Kraken2-detected respiratory pathogen. Bacterial (orange) and viral (red) pathogens NMHI distribution with 0.2 global model threshold for separating healthy vs unhealthy. The solid lines on ridge indicate medians.

## Discussion

This study establishes nasopharyngeal community state types as the organizing structure of the nasopharyngeal microbiome and demonstrates that failure to account for NPCST architecture explains contradictory findings that have impeded the field for over a decade. Six reproducible NPCSTs explain 52% of community variance and are associated with intrinsic diversity baselines largely independent of disease status. By resolving this diversity paradox through NPCST-aware analysis, we provide a stratified approach for identifying reproducible disease markers, one that body-site specific microbiome studies can adapt.

The diversity paradox we describe echoes a broader lesson from vaginal microbiome research, where community state types are similarly essential for interpreting diversity-disease relationships [79]. However, the nasopharyngeal case is distinct: unlike the vaginal microbiome where low diversity indicates health and high diversity signals dysbiosis [47, 79, 80], nasopharyngeal diversity reflects community architecture independent of wellness. This finding cautions against translating site-specific diversity assumptions to other microbiome niches without systematic evaluation. Our mixed-effects models show that diversity-disease associations are explained almost entirely by differential NPCST representation across study cohorts, suggesting that comparative nasopharyngeal analysis benefits from NPCST stratification or adjustment.

Beyond resolving diversity artifacts, NPCST-aware differential abundance analysis reveals that respiratory infection operates through two complementary mechanisms of community disruption. ANCOM-BC2 with LOSO cross-validation across 10 SARS-CoV-2 cohorts identified 13 reproducible taxa (9 enriched, 4 depleted; directional consistency ≥ 90%), including enrichment of obligate anaerobes associated with lower respiratory and gut compartments (*Faecalibacterium, Lachnospiraceae, Prevotellaceae*) [9, 81, 82], while all four depleted taxa were NP-resident hub genera (*Anaerococcus*, *Peptoniphilus*, *Lawsonella*, *Staphylococcus*) [29, 63]. Importantly, NPCST adjustment attenuated these signals by only 1.5%, confirming them as genuine disease taxa independent of community architecture. Co-occurrence network analysis independently converged on the same biological conclusion: the *Peptoniphilus-Anaerococcus* hub pair forms structural backbones across all NPCSTs, and hub genera exhibited 2.71-fold greater disruption during disease than non-hub taxa. This convergence of compositional evidence establishes that SARS-CoV-2 does not just alter abundance profiles, it also destabilizes the anaerobic backbone that maintains community stability while simultaneously permitting colonization by non-resident anaerobes. These subdominant anaerobes, previously understudied due to cultivation challenges [60], persist as network hubs despite the aerobic nasopharyngeal environment [29, 63, 75]. This paradoxical persistence likely reflects protected anaerobic micro-niches within mucosal biofilms, where oxygen consumption by facultative organisms generates gradients permitting obligate anaerobe survival, and disruption of these niches during infection may explain their preferential depletion. Individual components of this pattern have been verified by independent studies: *Faecalibacterium* as a discriminating NP genus [83], *Lachnospiraceae*/*Prevotellaceae* enriched in SARS-CoV-2-positive NP samples [9], and *Anaerococcus* and *Peptoniphilus* as core but subdominant NP colonizers [29, 63].

Together, these findings describe a reproducible pattern of backbone disruption across 10 LOSO cohorts. Notably, disruption of these same anaerobic core members has been independently associated with chronic respiratory conditions including COPD [29]. The consistent depletion of backbone genera during SARS-CoV-2 infection (100% directional consistency across cohorts) positions backbone integrity as both a candidate mechanistic target and a potential biomarker for community vulnerability evaluation.

The Nasopharyngeal Microbiome Health Index (NMHI) translates these ecological insights into a clinical application that achieved robust, generalizable discrimination across diverse disease conditions and all six NPCSTs in external validation. Unlike differential abundance approaches that produce study-specific biomarkers, NMHI aggregates community-wide presence/absence patterns into a continuous wellness metric, a conceptual shift from binary disease classification toward microbiome health monitoring. In the external validation, NMHI maintained robust performance across all six NPCSTs (AUC: 0.848-0.953) and differentiated disease populations with large effect sizes (median Cohen’s d = 2.179), confirming that community-level assessment generalizes beyond training conditions. The necessity of this approach is underscored by a fundamental paradox in nasopharyngeal biology: four of six NPCST keystone genera (*Haemophilus, Streptococcus, Moraxella, Staphylococcus*) contain species that are established respiratory pathogens yet simultaneously define healthy community states. Although *Streptococcus* and *Haemophilus* have been identified as network hubs [24, 64], their dual role as keystones and pathogens has not been formally contextualized. Detecting these organisms is clinically ambiguous without community context. *Haemophilus* dominates NPCST IV in healthy carriers just as it dominates during otitis media[26], and our meta-analysis identified *Streptococcus* colonizes 95.28% of all samples regardless of disease status. This duality renders single-genus biomarkers incomplete in the nasopharyngeal space and explains why prior machine learning classifiers targeting individual taxa achieved only low-to-modest performance (AUC 0.59-0.82) [27, 30, 35–37]. NMHI avoids this limitation by capturing the ensemble community signal via binarization of the detection signals. The biological basis for its robustness across all six NPCSTs is revealed by the convergence of network and NMHI analyses. Seven of nine hub genera conserved across ≥4 NPCSTs were independently selected as NMHI features, and LASSO further resolved species-level duality within these hubs, discriminating health-promoting from disease-promoting members of the same genus (e.g., *Haemophilus* genus vs *H. influenzae*) regardless of community state. This confirms that community-level health assessment is feasible across the nasopharyngeal microbiome’s architectural heterogeneity, because NMHI’s aggregation of NP-resident taxa across community types dilutes the effect of any single high-abundance organism. This is particularly relevant for the nasopharynx, where low microbial biomass amplifies the relative influence of any single high-abundance taxon on community-level metrics.

This framework has several limitations. First, short-read 16S rRNA gene sequencing constrains taxonomic resolution, yet our NMHI analysis demonstrates that species-level distinctions within the *Veillonella* hub genus carry opposing health signals (e.g., *Veillonella tobetsuensis* health-promoting vs *Veillonella parvula* disease-associated in the nasopharyngeal samples). Full-length 16S sequencing [14] represents the most feasible next step for improving resolution, as shotgun metagenomic approaches remain challenging for nasopharyngeal samples where host content exceeds 80% [82]. Second, geographic bias toward North America and Asia, combined with SARS-CoV-2 overrepresentation may limit generalizability, and broader cohorts spanning underrepresented populations and additional respiratory conditions/diseases are needed. Third, seasonal dynamics [29] and antibiotic exposure effects require longitudinal investigation within the NPCST framework. Fourth, while our decontamination pipeline demonstrated robust signal preservation, certain removed environmental-origin taxa (e.g., *Nocardiaceae* [83]) may be clinically relevant in immunocompromised populations, and the contamination reference should be refined for targeted studies. Finally, robust batch-effect assessment requires matched disease and control samples within each study. Our discovery cohort, comprising remnant specimens submitted for clinical respiratory panel testing, contained only symptomatic patients and therefore lacked healthy controls from the same sequencing batch. Future validation studies incorporating matched healthy controls would enable within-batch differential abundance testing to independently confirm the identified microbial disease markers.

Our framework suggests possibilities for NPCST-targeted therapeutics. Specifically, NMHI provides a continuous wellness metric enabling graded risk stratification rather than binary designations confounded by obligate and opportunistic pathogen co-detection. The distinct disease susceptibilities across NPCSTs, particularly NPCST VI’s association with bacterial infection, and the central role of anaerobic network hubs suggest opportunities for precision interventions. As millions of nasopharyngeal swabs are collected annually for respiratory diagnostics, integrating microbiome assessment alongside pathogen detection points toward more in-depth respiratory health evaluation. As community structure differs by body site, future work should apply this similar stratified framework elsewhere to test, rather than assume, whether these findings transfer and to clarify genuine site-specific microbial biology.

## Ethics approval and consent to participate

The use of residual de-identified samples for this study was determined as not human subject research requiring Institutional Review Board (IRB) review according to WIRB-Copernicus Group (WCG) IRB. All samples were residual clinical materials originally collected for routine diagnostic testing and were fully de-identified prior to investigator use; investigators had no interaction or intervention with the individuals from whom the specimens originated. Informed consent to participate was not required on this basis. All work was performed in accordance with the Declaration of Helsinki.

## Availability of data and materials

The data that support the findings of this study are available from multiple sources. Primary data analyzed in this meta-analysis are openly available in the source publications and corresponding NCBI BioProjects as listed in **Table 1**. The discovery set 16S data can be found on BioProject PRJNA1509995. The NPCST classifier machine learning models are available in Zenodo (DOI: 10.5281/zenodo.17068997). Additional scripts used as part of Labcorp’s internal bioinformatics workflow are not publicly available due to institutional data-governance policies. All analytical approaches and computational methods applied are fully described in the **Supplementary File**, and further methodological detail is available from the corresponding author upon reasonable request.

## Author Contributions

K.S. performed the validation and visualization. K.S., H.B., and L.I. conceived this study. K.S., H.B., M.B., Q.Z., M.T., and A.H. performed data curation and contributed resources.

K.S. and H.B. developed the methodology and conducted formal analysis. K.S. and L.I. provided supervision and project administration. C.I. and S.L. acquired funding. K.S. wrote the original draft, with all authors contributing to manuscript review and editing. All authors read and approved the final manuscript.

## Acknowledgements

We would like to thank the U.S. Centers for Disease Control and Prevention (CDC) for their guidance and expertise throughout the REaSoN project.

---

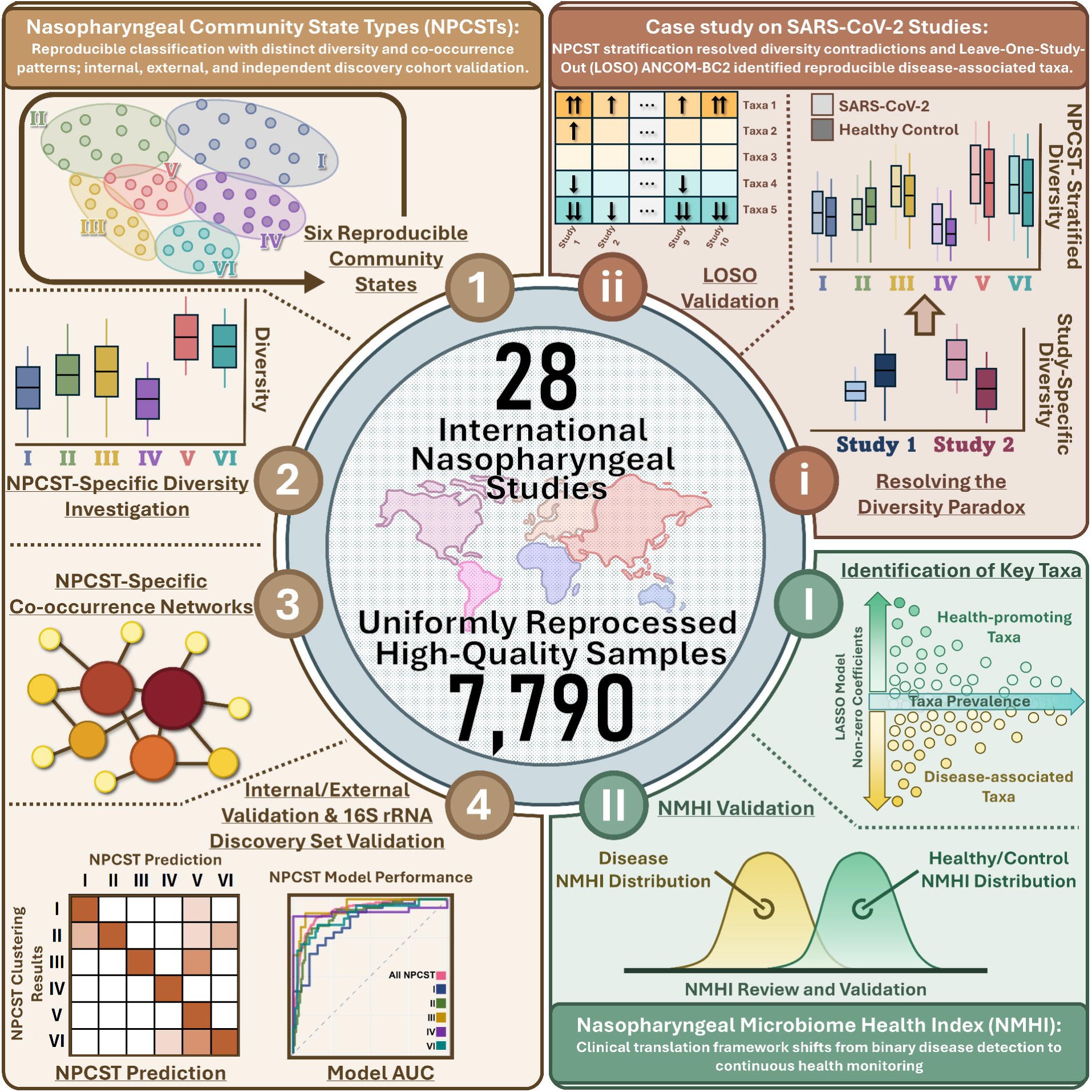

