## Supplementary File for "Reproducible Community States Resolve the Nasopharyngeal Diversity Paradox"

### Introduction

The following documents include the supplementary methods and results for many of the analyses that were conducted to support the main publication. If the methods were detailed in the main manuscript, we refrained from including the duplicated text in this document.

### Methods

#### Large language model-assisted background contamination screening

We employed Claude [1] (Anthropic) to systematically evaluate bacterial families for potential contamination signatures. Each SILVA-annotated family underwent independent triplicate assessment using a standardized prompt designed to identify taxa inconsistent with nasopharyngeal ecology (**Figure 2A**). The LLM evaluated each family for likelihood of representing reagent contamination ("kitome")[2, 3], environmental sources (water, soil, laboratory), or legitimate nasopharyngeal colonizers. Consensus classification required agreement across all three independent queries, with discordant cases flagged for manual expert review. A microbiome specialist with expertise in respiratory tract ecology performed final adjudication of ambiguous classifications, incorporating published nasopharyngeal microbiome literature and contamination databases. This human-in-the-loop approach ensured that AI-assisted screening augmented rather than replaced expert taxonomic knowledge, consistent with best practices for AI implementation in microbiome research[4].

#### Prompt for contamination assessment

"You are a nasopharyngeal microbiome expert tasked with identifying potential contaminants in 16S rRNA sequencing data. Evaluate the provided bacterial families for consistency with genuine nasopharyngeal colonization versus likely contamination from reagents, water sources, laboratory environments, or sample processing artifacts. Consider typical nasopharyngeal ecology, human commensal flora, and known contamination patterns in low-biomass samples.

Return results in CSV format with columns:

1. Bacterial Family (string)
2. Contaminant (boolean: TRUE/FALSE)
3. Source (string: Reagent/Water/Soil/Laboratory/Environmental/Not_applicable)

#### 16S data processing

Raw sequencing data processing was performed using R v4.1.1[5]. Hypervariable region (HVR) targets and primer presence/absence were first identified by constructing a comprehensive list of possible HVR primer pairs and their maximum expected insert sizes (**Tables S1** and **S2**). A kmer hash of each possible HVR configuration was built by in silico amplifying various HVRs from the SILVA v138.2 database [6] using Mash with kmer size 31[7]. When kmer hashing could not distinguish HVRs but PCR primers were present, HVR configuration was inferred directly from the PCR primers (**Table S1**). This process independently verified the HVRs reported in corresponding source publications (**Table 1**). When PCR primers were detected, reads were reoriented to the same strand using an in-house R function utilizing the ShortRead [8] R package.

Quality filtering and trimming were performed using the DADA2 v1.22.0 [9] filterAndTrim function. Parameters were optimized using an in-house R function that accounted for read lengths and quality profiles, presence/absence of PCR primers, and maximum expected hypervariable region insert sizes (**Table S2**). Quality-filtered trimmed reads underwent standard DADA2 processing including denoising and merging to generate amplicon sequence variants (ASVs). Singletons and chimeras were removed during ASV filtering.

Taxonomic classification utilized SILVA v138.2 database [1] training data formatted for DADA2 (obtained from Zenodo, DOI: 10.5281/zenodo.14169026). The DADA2 assignTaxonomy function was applied with minBoot=80 using silva_nr99_v138.2_toGenus_trainset.fa.gz, followed by the addSpecies function with default parameters using silva_v138.2_assignSpecies.fa.gz. ASVs lacking at least family-rank classification were excluded from analysis. Final ASVs were aggregated at the lowest assigned taxonomic rank to generate count matrices for each study.

Beyond the contamination removal procedures described in the main manuscript, we implemented stringent quality control criteria at both sample and study levels. Individual samples required a minimum of 5,000 reads successfully mapped to family-level taxonomy following contaminant family removal to ensure adequate sequencing depth for reliable taxonomic profiling. To maintain dataset consistency for meta-analysis, we further required that studies retain at least 50% of samples passing these quality thresholds for inclusion in the final analysis. This two-tiered filtering strategy eliminated both low-quality samples and studies with systematic technical failures while preserving comparability that powered our study.

#### Phylogenetic analysis

Since the meta-analysis spanned multiple hypervariable regions (Table 1), V3-V4 ASVs were selected for phylogenetic analysis based on the taxonomic representation assessment described in the taxonomic resolution comparison above. ASVs were clustered using CD-HIT v4.8.1 [11] at 99% identity, with representative ASVs selected by prioritizing unique and highly resolved taxonomic assignments. Specifically, when multiple ASVs shared identical taxonomy at the genus or species level, the representative ASV was selected based on (1) assignment to a unique taxonomic lineage not represented by other cluster representatives, and (2) the highest taxonomic resolution (species > genus > family). Following clustering, ASVs required detection in a minimum of two studies to ensure cross-study reproducibility. A multiple sequence alignment was generated using Clustal Omega v1.2.4 [12], and a Jukes-Cantor distance-optimized maximum likelihood tree was constructed using phangorn v2.8.1 [13] in R and visualized with ggtree v3.2.0 [14].

#### NPCST classification comparison between the before and after-background decontamination

Cluster preservation was evaluated using the Adjusted Rand Index (ARI) to quantify agreement between before and after background decontamination cluster assignments, with bootstrap resampling (n = 100) generating 95% confidence intervals. This evaluation used only the 7,790 high-quality samples retained in the final dataset after background decontamination. Internal cluster validity was assessed through silhouette analysis, which measured within-cluster cohesion relative to separation from neighboring clusters. Silhouette coefficients were calculated using Bray-Curtis dissimilarity matrices for k = 6 clusters derived from Ward's hierarchical clustering, enabling direct comparison of clustering quality between before and after background decontamination datasets.

To validate preservation of microbiome community relationships following decontamination, we employed complementary ordination-based approaches using

Procrustes analysis and Mantel tests on Bray-Curtis distance matrices. Procrustes analysis optimally rotated and scaled principal coordinate analysis (PCoA) ordinations to maximize alignment between before and after background decontamination datasets, yielding a correlation coefficient and $M^2$ statistic with 999 permutations. The Mantel test independently evaluated the correlation between pairwise sample distances in both distance matrices using Pearson correlation coefficients with 999 permutations.

#### Taxonomic resolution comparison between V3-V4 and V4

To determine the appropriate taxonomic rank for meta-analysis, we compared taxonomic resolution between genus and species levels using only studies employing V3-V4 or V4 hypervariable regions, as other regions were rare in our dataset and did not warrant comparison. We performed rarefaction analysis to assess taxonomic saturation at both ranks, calculated the cumulative relative abundance of taxa shared between V3-V4 and V4 regions, and quantified the proportion of sequences that could not be classified at each taxonomic level. Taxa with relative abundance below 0.01% were excluded from all analyses to minimize noise from rare sequences.

#### Leave-one-study-out (LOSO) NPCST investigation

To assess the reproducibility and stability of identified NPCSTs, we implemented a leave-one-study-out cross-validation approach across all 28 studies. For each iteration, one study was systematically removed, and hierarchical clustering was performed on the remaining 27 studies using Bray-Curtis dissimilarity at the genus level followed by Ward linkage, with cluster assignments tested for k values ranging from 4 to 12. The complete 28-study dataset served as ground truth, and generated clusters were matched to reference NPCSTs using a greedy assignment algorithm based on contingency table analysis, prioritizing clusters with the highest intersection-to-union ratios.

Clustering stability was quantified using three metrics: (1) Adjusted Rand Index (ARI) to measure overall clustering agreement corrected for chance, (2) Mean Jaccard Index to assess average per-cluster similarity between predicted and ground truth assignments, and (3) Overall Accuracy to calculate the proportion of samples correctly assigned to their corresponding ground truth NPCST after optimal cluster matching.

#### Unsupervised learning based definition of the rare biosphere (ulrb)

To objectively define abundance categories within each CST, we applied the ulrb[15] method as described by Pascoal et al. Following the recommended approach, we employed the default tri-categorization framework (k=3) to classify genera into "Abundant", "Undetermined", and "Rare" categories within each sample. The quality of clustering was evaluated using Silhouette scores, with scores >0.5 indicating reasonable to strong cluster structure across all CSTs. Abundant genera within each CST were characterized by their median relative abundance, detection prevalence across samples, and clustering quality metrics.

#### Machine learning approach for validating NPCST classification

We developed and validated a comprehensive machine learning framework to classify nasopharyngeal swab samples into six previously defined NPCST categories using relative abundance data from 626 genera across 7,790 samples spanning 28 independent studies. We validated this model using 28 studies and further evaluated its performance on two external datasets. Each method underwent hyperparameter optimization and evaluation through 100 iterations of 5-fold cross-validation, with performance assessed using accuracy, precision, recall, and F1-score metrics to ensure comprehensive evaluation across all NPCST categories. To ensure robust model generalization across diverse study populations, we implemented a stratified cross-validation strategy that maintained balanced distribution of both target NPCST classifications and source studies (BioProjects) origins within each fold, thereby preventing potential batch effects from influencing model performance and enhancing the generalizability of the final model.

For machine learning model selection, we focused on algorithms widely used in the microbiome field and methodologically distinct approaches across tree-based, regression-based, and kernel-based methods. The selected models included Random Forest (randomForest [16] v4.7-1.2) with hyperparameters including mtry values ranging across different numbers of genera and ntree values of 50, 100, 200, 500, and 1,000 trees; Ridge, LASSO, and elastic net regression (glmnet[17] v4.1-9) with regularization parameter λ optimized through 5-fold cross-validation and alpha fixed at 0 (Ridge), 0.1–0.9 (Elastic Net), and 1 (LASSO); and Support Vector Machine (e1071[18] v1.7-16) with radial basis function kernel, cost parameters ranging from 0.1 to 100, and gamma parameters from 0.001 to 1.0. For each of the iterations and individual 5-fold cross validation, each of these hyperparameter was examined and the best one is recorded.

During evaluation, we removed LASSO regression from analysis because many 5-fold cross-validation sets failed to converge, resulting in over 30% missing data points. We calculated performance metrics using the caret[19] package (v7.0-1), employing balanced accuracy to account for potential class imbalances, where overall accuracy represented the macro-averaged balanced accuracy across all classes and per-class accuracy corresponded to individual class balanced accuracy. Additionally, we tracked Random Forest feature importance through mean decrease in Gini impurity and mean decrease in accuracy.

#### NPCST classification model deployment

The final SVM and Random Forest models were independently trained on the complete dataset of 626 genera across 7,790 samples from 28 independent studies. We developed customized functions to enable future users to apply these models to new datasets. The deployment pipeline validates genera naming conventions before performing NPCST classifications and generates confidence scores for both SVM and Random Forest predictions. Complete implementation instructions are provided in the Zenodo repository (DOI: 10.5281/zenodo.17068997).

#### External Validation dataset

Two additional BioProjects were selected from publications released after the initial extraction cutoff for NPCST classifier validation: PRJNA981220 (SARS-CoV-2, n = 40) and PRJNA914884 (respiratory syncytial virus (RSV), n = 1,537). These were processed using identical protocols, yielding 1,577 samples with NPCST assignments from Random Forest classification. NMHI external validation used an independent set of 699 longitudinal and cross-sectional samples not included in model training.

#### External validation of the NPCST prediction model

For external validation evaluations, input data underwent the nasopharyngeal-specific background removal protocol followed by data validation and NPCST prediction according to the deployed classification guide available in the Zenodo repository. We established ground-truth classifications for these external validation samples using the same Bray-Curtis dissimilarity followed by Ward linkage methodology, combining the 28 original studies with the 2 external validation studies or our own discovery set, then selected the top 6 NPCSTs. We performed ROC calculations using the pROC [20] package (v1.18.5) in R.

#### Independent PCR Panel

We processed 124 specimens via Seegene Novaplex™ multiplex RT-PCR (Respiratory 1a, 2, 3 & PneumoBacter panels, 26 targets) to characterize co-detection patterns. Individual-level SeeGene Ct data and metatranscriptomics results will be reported in another publication focused on total RNA-seq respiratory pathogen surveillance; summary co-detection statistics are presented herein. The four panels are:

- **Novaplex™ Respiratory Panel 1A** (7 targets): Influenza A virus (with H1, H1pdm09, H3 subtyping), Influenza B virus, RSV A, RSV B
- **Novaplex™ Respiratory Panel 2** (7 targets): Adenovirus, Enterovirus, Metapneumovirus, Parainfluenza virus 1–4

- **Novaplex™ Respiratory Panel 3** (5 targets): Bocavirus 1/2/3/4, Rhinovirus A/B/C, Coronavirus NL63, Coronavirus 229E, Coronavirus OC43
- **Novaplex™ PneumoBacter Assay** (7 targets): *Bordetella pertussis*, *B. parapertussis*, *Chlamydophila pneumoniae*, *Haemophilus influenzae*, *Legionella pneumophila*, *Mycoplasma pneumoniae*, *Streptococcus pneumoniae*

Together, these four panels interrogate 26 distinct respiratory pathogen targets spanning the major viral and atypical bacterial agents of respiratory tract infection. Samples were classified as positive for a given target when amplification curves crossed the manufacturer-specified threshold, with Ct values recorded for all positive detections.

#### Co-detection analysis

For co-detection quantification, internal control (IC) amplifications were excluded. Influenza A subtypes (H1, H1pdm09, H3) were consolidated under a single "Influenza A" designation, as subtype-level co-detection with the parent target reflects assay design rather than biological co-detection. Each sample was then classified by co-detection category: single pathogen (one target detected), viral–bacterial co-detection (at least one viral and one bacterial target), multi-viral co-detection (two or more distinct viral targets), or multi-bacterial co-detection (two or more bacterial targets). The individual-level Seegene Ct data are not released in this publication; summary co-detection statistics and figures are provided herein to characterize the poly-microbial landscape that motivated the community-level NMHI approach.

#### Nasopharyngeal microbiome health index (NMHI)

Adapted from Chang et al.'s GMWI2 methodology[21], we developed the Nasopharyngeal Microbiome Health Index (NMHI) using LASSO-penalized logistic regression (glmnet R package v4.1) with balanced class weights to address sample size imbalances. We calculated class weights as ($w_{healthy} = 0.5/(n_{healthy}/n_{total})$, $w_{disease} = 0.5/(n_{disease}/n_{total})$), ensuring equal class contribution regardless of imbalance. From 5,435 cross-sectional nasopharyngeal samples, we constructed binary presence/absence matrices using a 0.01% relative abundance threshold (present=1, absent=0).

We implemented four binary classification models to four distinct models: (1) healthy controls versus combined diseased samples with all-taxa (All-taxa All-Conditions), (2) healthy controls versus viral infections with all-taxa (All-taxa Viral Infection), (3) healthy controls versus combined diseased samples with genus-only taxa (Genus-only All-Conditions), and (4) healthy controls versus viral infections with genus-only taxa (Genus-only Viral Infection). This design enabled assessment of both taxonomic granularity and disease specificity effects on model performance.

We evaluated seven taxonomic configurations (all taxa combined, phylum, class, order, family, genus, and species), excluding unclassified reads to ensure interpretability of results. Model development and validation proceeded through five stages:

**Stage 1: Leave-One-NPCST-Out Cross-Validation for Lambda Selection:** We implemented LONO cross-validation to establish stable lambda values, leveraging the biological distinctiveness of NPCSTs and their differential disease susceptibilities. This approach systematically held out each NPCST (I-VI) as a test set while training on the remaining five, ensuring generalization across biologically meaningful community states rather than technical batch effects. Given that NPCSTs explained substantially more variance than study effects (52.19% vs. 13.06%), this strategy provided robust parameter selection. We tested selective lambda values ranging from 0.0001 to 0.03, selecting optimal values based on maximum AUC aggregated across all six held-out NPCST test sets.

**Stage 2: Model Performance Evaluation with Prevalence Thresholds:** Using optimal lambda values from Stage 1, we performed both LONO and 10-fold cross-validation at five prevalence thresholds (0%, 1%, 5%, 10%, 20%) to assess model robustness. The LONO is run only once for the cross-validation and the 10-fold cross-validation repeated 10 times with reproducible seed numbers (seeds 1-10) to ensure reproducibility while capturing variability. We maintained strict train-test separation within each fold to prevent data leakage, confining lambda selection exclusively to training partitions. The NMHI score was calculated as the sum of products between coefficients and binary presence values, where positive coefficients indicated health-associated taxa and negative coefficients indicated disease-associated taxa. We assessed model performance using both AUC and balanced accuracy metrics to evaluate discriminatory power and classification performance.

**Stage 3: Final Model Training:** Using the selected 5% prevalence threshold (based on optimal performance-interpretability trade-off), we trained final models on the complete dataset with optimal lambda values. During this stage, we extracted all non-zero coefficients and intercepts from each model, enabling NMHI score calculation as the sum of the intercept and products of coefficients with presence/absence values for each sample's taxa. To identify key microbial markers, we analyzed taxa with absolute coefficients ≥0.5 and performed comparative abundance analysis across control and disease groups. We visualized abundance distributions using boxplots and assessed statistical significance using Wilcoxon rank-sum tests with FDR correction for multiple testing comparisons.

**Stage 4: NMHI Threshold Optimization:** Using models trained on the complete dataset, we optimized classification thresholds to distinguish healthy from diseased samples. We evaluated both global (across all NPCSTs) and NPCST-specific thresholds ranging from -5 to 5 (at 0.1 increments), selecting optimal values based on maximum balanced accuracy for each model configuration. We calculated Cohen's d effect sizes to quantify the magnitude of separation between healthy and diseased populations, providing a standardized measure of discriminatory power independent of sample size. We also performed Wilcoxon rank-sum tests with FDR correction for multiple testing to statistically compare disease and control samples for both global and NPCST-specific models.

**Stage 5: External Validation:** External validation utilized both cross-sectional and longitudinal samples (n=699 total), treating each sampling point as independent given the transient nature of nasopharyngeal microbiome communities during infection. This approach tests the model's ability to distinguish disease states regardless of sampling design, reflecting real-world diagnostic applications where single-timepoint sampling is standard. To ensure valid assessment, we removed samples with ambiguous disease classifications (e.g., pneumococcal carriers, emergency room volunteers) and samples not consistently obtained during symptomatic disease periods. The validation cohort comprised healthy family members with recurrent respiratory tract infections (n=265), varying SARS-CoV-2 severity (n=398), lower respiratory tract infections (n=5), and non-SARS-CoV-2 critically ill patients (n=31). We applied final model coefficients to calculate NMHI scores and generate predictions for these previously unseen samples, evaluating performance using ROC (Receiver Operating Characteristic) curve analysis and balanced accuracy metrics with both global and NPCST-specific thresholds. The NPCST classification was based on the random forest model we provided in the earlier section. For NPCST-specific validation, we excluded NPCST V from AUC analysis when all samples belonged to a single class (disease), as meaningful discrimination requires representation of both classes.

#### Statistical methods

All statistical analyses were performed in R (v4.2.2) [5]. Alpha diversity metrics (genus richness, Shannon diversity, Inverse Simpson) were calculated after rarefying to 5,000 reads using vegan [22] package to standardize sequencing depth and stabilize diversity estimates across the 28 pooled studies' heterogeneous sequencing depths. Decontamination validation employed Procrustes analysis and Mantel tests (999 permutations). Group comparisons used Wilcoxon rank-sum tests with Benjamini-Hochberg correction. Effect sizes were reported as Cohen's d (continuous), odds ratios with 95% CI (categorical), and epsilon-squared or Cramer's V (NPCST-demographic associations). Model performance was assessed using AUC with 95% confidence intervals and balanced accuracy at optimized thresholds.

### Results

#### Validation of signal integrity following background decontamination

Following implementation of our three-stage decontamination pipeline (**Figure 2A**), we evaluated quality control metrics across all 28 studies to validate background removal effectiveness. The sigmoid distribution patterns observed across studies demonstrated that most samples retained robust true signal abundance (>80% cumulative relative abundance) with >5,000 reads after contamination removal, exhibiting minimal background interference (**Figure S1**). Of 8,314 total samples, 7,986 (96.1%) successfully exceeded the 5,000 true-signal read threshold, confirming that our decontamination approach preserves sufficient sequencing depth for downstream analyses. This consistent retention pattern across diverse studies validates our pipeline's capacity to eliminate spurious signals while maintaining biological integrity. The final dataset comprised 7,790 samples after additional quality control for complete disease/health status annotation and exclusion of rare positive and negative control samples. After applying the same blacklist background removal protocol, we revealed only three novel genera (*Tersicoccus*, *Bact-08*, *Eoetvoesia*) absent from our training data, all at minimal abundances around 0.02%, demonstrating comprehensive capture of the core nasopharyngeal microbiome across diverse populations and confirming model applicability to new cohorts.

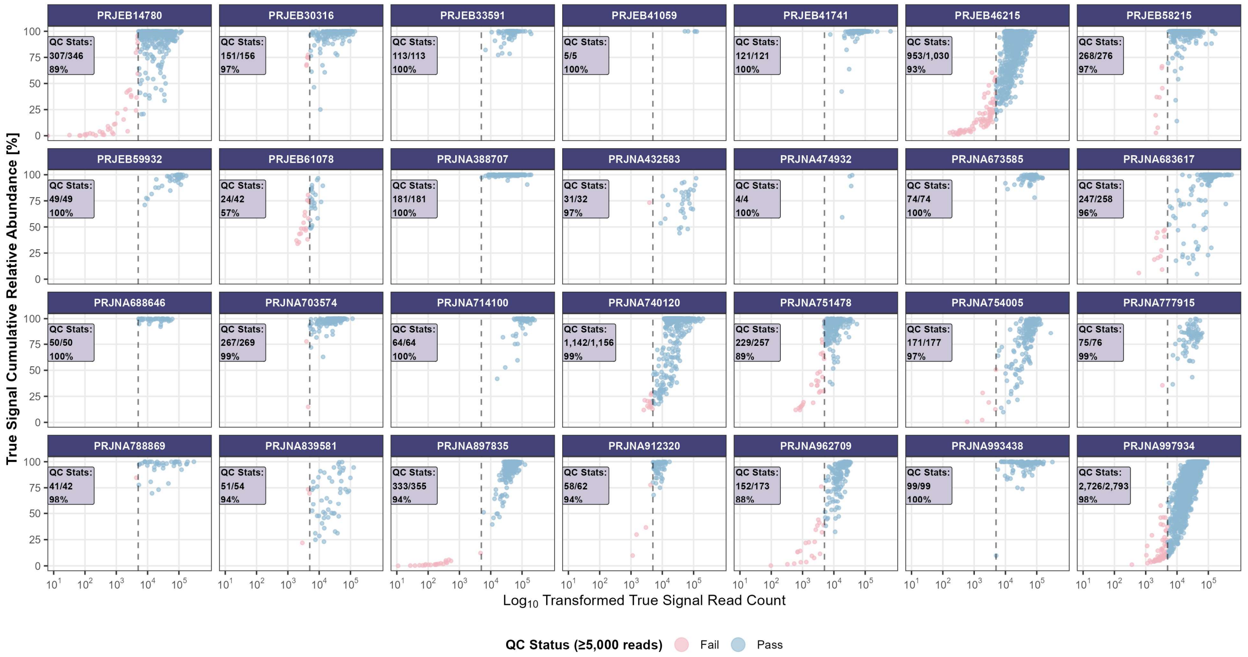


**Figure S1. Study-specific quality control assessment.** Quality control evaluation of nasopharyngeal microbiome samples across 28 retained studies, displaying the relationship between $\log_{10}$-transformed true signal read counts and cumulative relative abundance following background removal. Each panel presents study-specific retention statistics (passed/total samples and percentage) based on the 5,000-read threshold criterion, with blue points representing QC-passed samples and pink points indicating failed samples.

#### Taxonomic resolution comparison between V3-V4 and V4

Rarefaction analysis revealed that the majority of samples reached diversity plateaus before 5,000 reads for both genus and species levels, confirming the adequacy of our quality control parameters (**Figure S2A-B**). Although selected samples with higher microbial richness required over 10,000 reads for complete saturation, genus-level rarefaction curves consistently plateaued earlier than species-level curves across all samples. Evaluation of cumulative relative abundance of shared taxa between V3-V4 and V4 regions identified 485 shared genera and 887 shared species (**Figure S2C**). Notably, nearly all V4-identified taxa were present within the V3-V4 dataset, while V3-V4 contained additional unique taxa, and expected pattern given the inclusion of the V3 region. Genus-level classification demonstrated significantly better consistency between regions, with virtually all taxonomic assignments shared between V3-V4 and V4, enabling harmonized analyses independent of hypervariable region selection.

Analysis of unassigned sequences revealed substantial differences in classification success between taxonomic levels, regardless of hypervariable region (**Figure S2D**). Species-level classification failed for a median of 75.9% (V3-V4) and 84.7% (V4) of reads, with high variability across studies (SD = 25.0% and 18.1%, respectively). Nearly all studies contained samples with >50% species-level assignment failure, creating severe resolution imbalances that would compromise downstream analyses. In contrast, genus-level classification maintained robust performance with median unassigned proportions of only 0.4% (V3-V4) and 0.6% (V4). These findings demonstrate that genus-level classification provides the taxonomic resolution necessary for reliable meta-analysis across heterogeneous nasopharyngeal microbiome studies from 16S data.

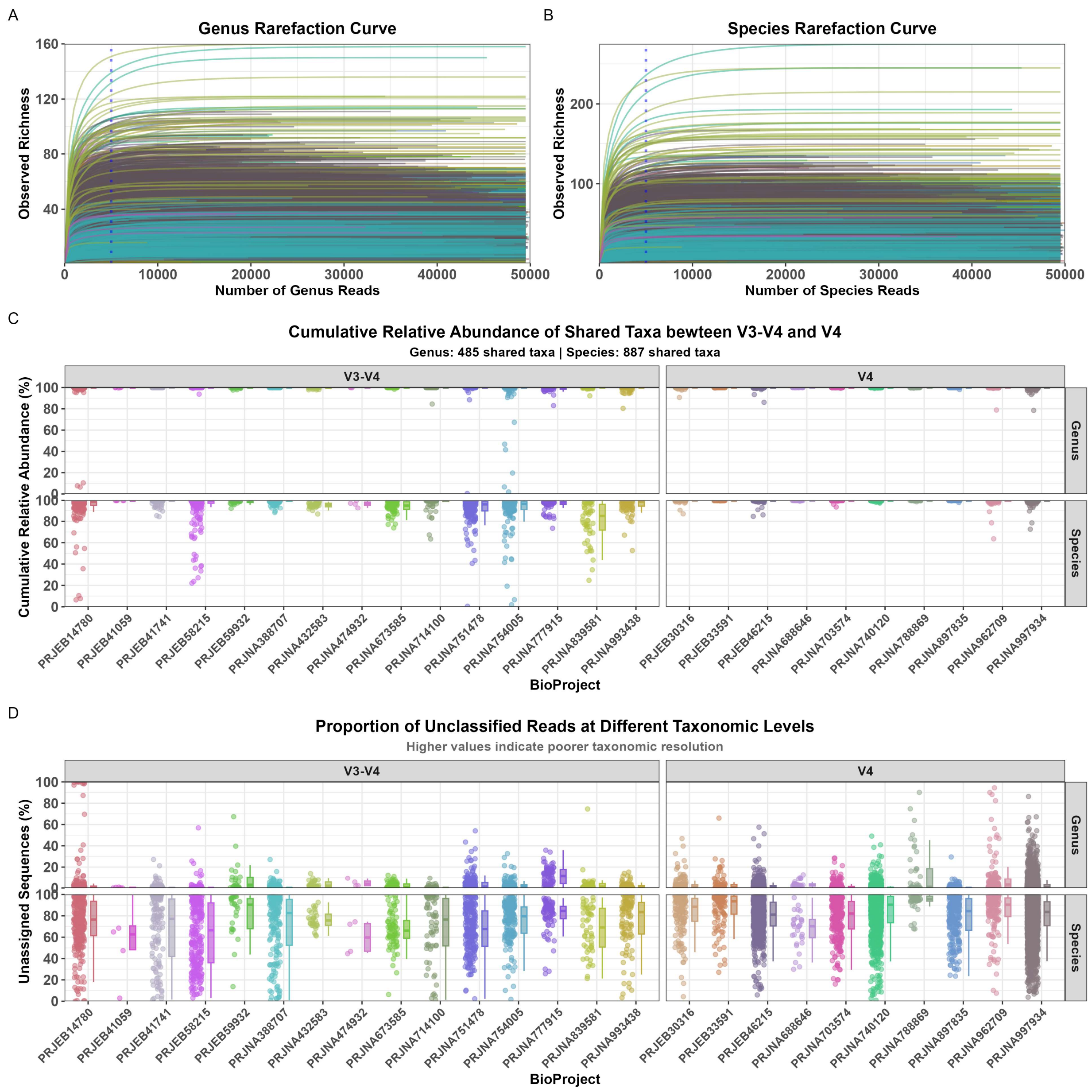


**Figure S2. Taxonomic resolution comparison between genus and species levels for V3-V4 and V4 16S rRNA gene regions in nasopharyngeal microbiome meta-analysis. A-B**. Rarefaction curves demonstrate that genus-level diversity approaches saturation while species-level richness continues to increase without plateauing, with a vertical blue dotted line indicating the 5,000 read threshold. **C**. Cumulative relative abundance of shared taxonomic features between V3-V4 and V4 regions reveals that genus-level classification captures substantially higher proportions of the microbial community (485 shared genera) compared to species-level classification (887 shared species), stratified by hypervariable region and BioProject. **D**. Proportion of unclassified sequences shows consistently higher assignment failure rates at species level compared to genus level across both V3-V4 and V4 regions, indicating poor species-level resolution reliability. Colors throughout all panels represent distinct BioProjects included in the meta-analysis.

#### Comparison of microbial community structure before and after background decontamination

Comprehensive validation analyses confirmed that our decontamination pipeline preserved biological signal integrity while enhancing data quality. Cluster assignments demonstrated high stability, with 6,327 of 7,790 samples (81.2%) maintaining their original nasopharyngeal community state type classification and moderate-to-strong agreement between before and after background decontamination datasets (ARI = 0.605, 95% CI: 0.592–0.617). Clustering quality improved substantially by 18.8%, increasing from 0.245 to 0.291, with cluster and group transitions visualized in **Figure S3A–B**. Procrustes analysis revealed near-perfect preservation of sample relationships (correlation = 0.973, $M^2$ = 0.05, $p < 0.001$), corroborated by Mantel test results showing exceptionally strong correlation between distance matrices ($r = 0.971$, $p < 0.001$). These convergent lines of evidence demonstrate that removing 1,810 contaminating genera (reducing the dataset from 2,436 to 626 genera) enhanced detection of genuine nasopharyngeal microbiome patterns without distorting underlying biological relationships. Principal coordinate analysis further validated this preservation, with the first two dimensions each exhibiting approximately 2% increased variance explained after background decontamination, indicating improved sample separation (**Figure S3C–D**). The reduction in study-associated variance from 13.97% to 13.06% ($R^2$ from PERMANOVA) represents meaningful mitigation of batch effects. Sparsity decreased from 97.89% to 95.53% following decontamination, as removing 1,810 predominantly sparse contaminating genera eliminated approximately 14 million data points (mostly zeros) while resulting in a denser matrix with 2.36 percentage points fewer zeros, thereby improving data quality for downstream analyses.

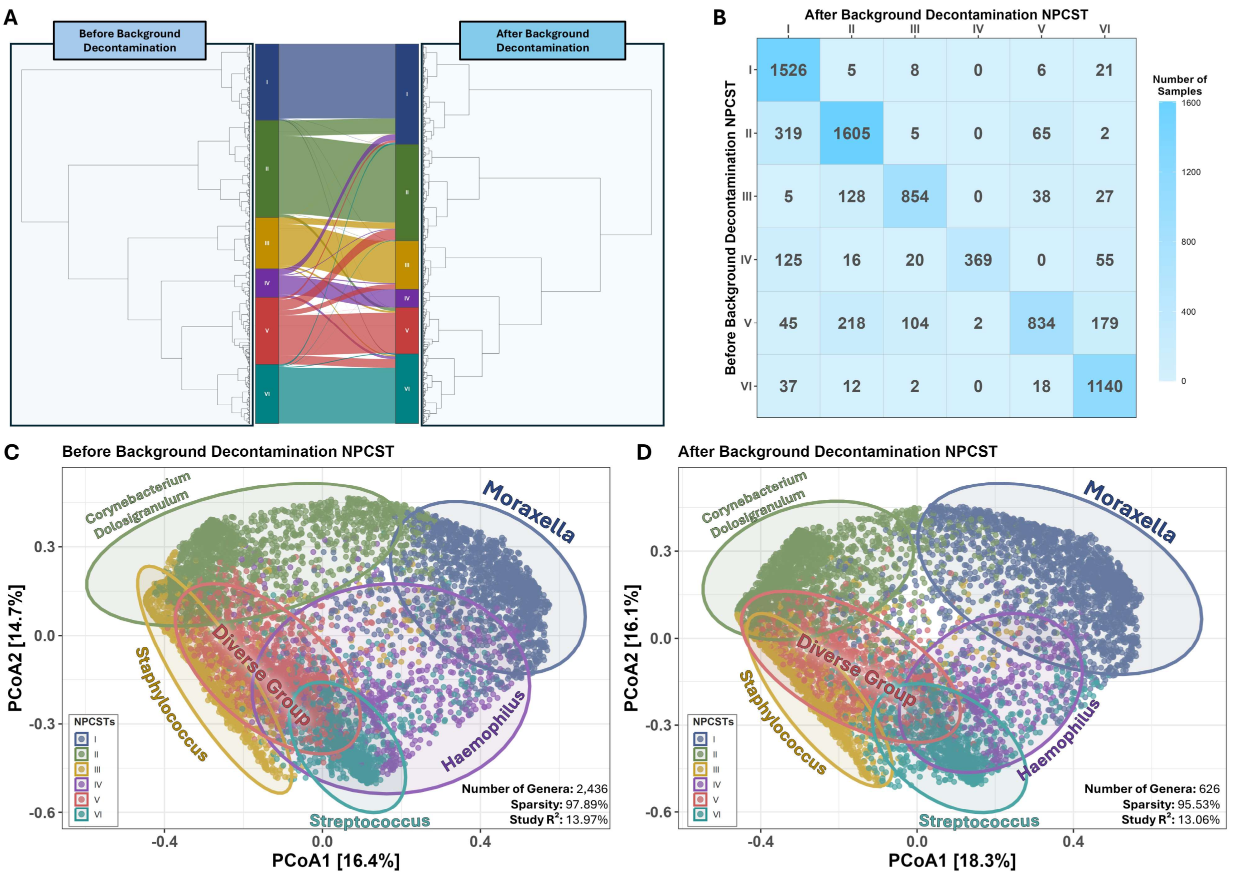


| | I | II | III | IV | V | VI |
|---|---|---|---|---|---|---|
| I | 1526 | 5 | 8 | 0 | 6 | 21 |
| II | 319 | 1605 | 5 | 0 | 65 | 2 |
| III | 5 | 128 | 854 | 0 | 38 | 27 |
| IV | 125 | 16 | 20 | 369 | 0 | 55 |
| V | 45 | 218 | 104 | 2 | 834 | 179 |
| VI | 37 | 12 | 2 | 0 | 18 | 1140 |

**Figure S3. Comparison of microbial NPCST structure before and after background decontamination. A.** Hierarchical clustering dendrograms displaying six NPCSTs identified in before and after background decontamination datasets, with an alluvial plot illustrating sample transitions between corresponding clusters. **B.** Confusion matrix quantifying the redistribution of samples across NPCSTs from before background decontamination (x-axis) to after background decontamination (y-axis) stages. **C&D.** Principal coordinate analysis (PCoA) ordination of the two PCoA dimensions before and after decontamination datasets, respectively. Ecological metrics including the number of retained genera, matrix sparsity (genera × samples), and variance explained ($R^2$) from PERMANOVA analysis are displayed in the lower right corner of each panel.

#### NPCST-specific cumulative relative abundance of top 14 families

We examined the NPCST-specific median cumulative abundance patterns (**Figure S4**). The top six families elevated most samples in NPCSTs I-IV to achieve 80-90% cumulative relative abundance. In contrast, the more diverse NPCSTs V and VI demonstrated slower abundance progression. Collectively, the top 14 families explained a median of at least 90% of total relative abundance across all samples and NPCSTs, confirming that these 14 families (comprising 161 genera and 513 ASVs) represent the core nasopharyngeal bacterial composition.

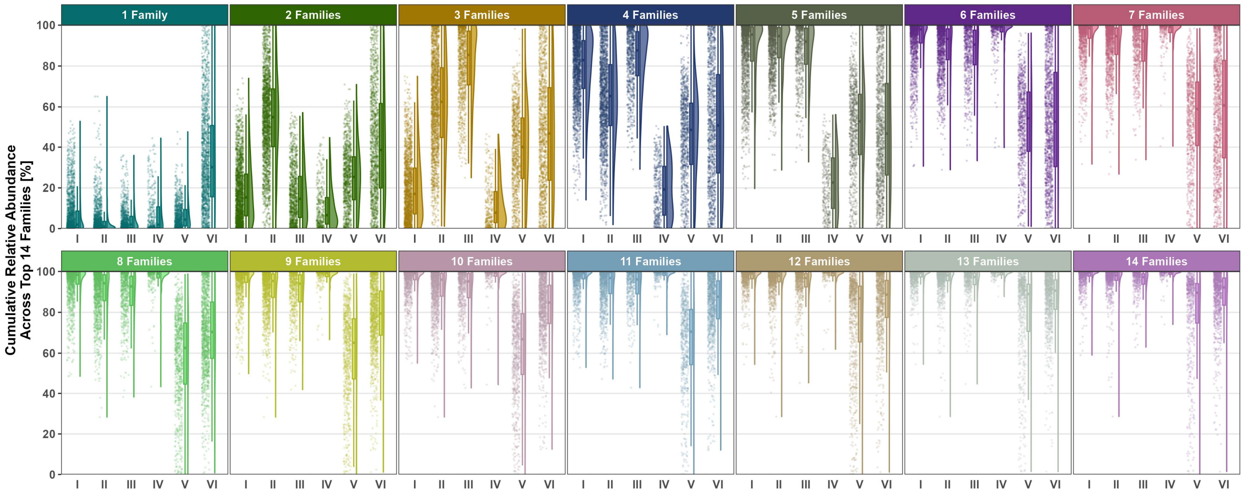


**Figure S4. NPCST-specific cumulative relative abundance of top 14 families.** Cumulative relative abundance distributions across the top 14 most prevalent families after background decontamination, with each panel representing sequential accumulation of relative abundance from the highest-ranked family through progressively lower-ranked families (left to right), demonstrating sample-level variability within each cumulative stage across six NPCSTs.

#### LOSO validations

Leave-one-study-out cross-validation analysis identified 6 NPCSTs as the optimal clustering solution, demonstrating robust stability across all evaluated metrics (**Figure S5**). At k=6, the Adjusted Rand Index achieved 0.711 and the mean Jaccard Index reached 0.75, indicating strong clustering agreement and high per-cluster similarity that substantially exceeded random performance. Overall median accuracy attained 0.867, demonstrating that 87% of samples were correctly assigned to their corresponding ground truth NPCSTs during cross-validation. These converging stability metrics collectively support the selection of 6 NPCSTs as the most reproducible and biologically meaningful clustering structure in the nasopharyngeal microbiome data.

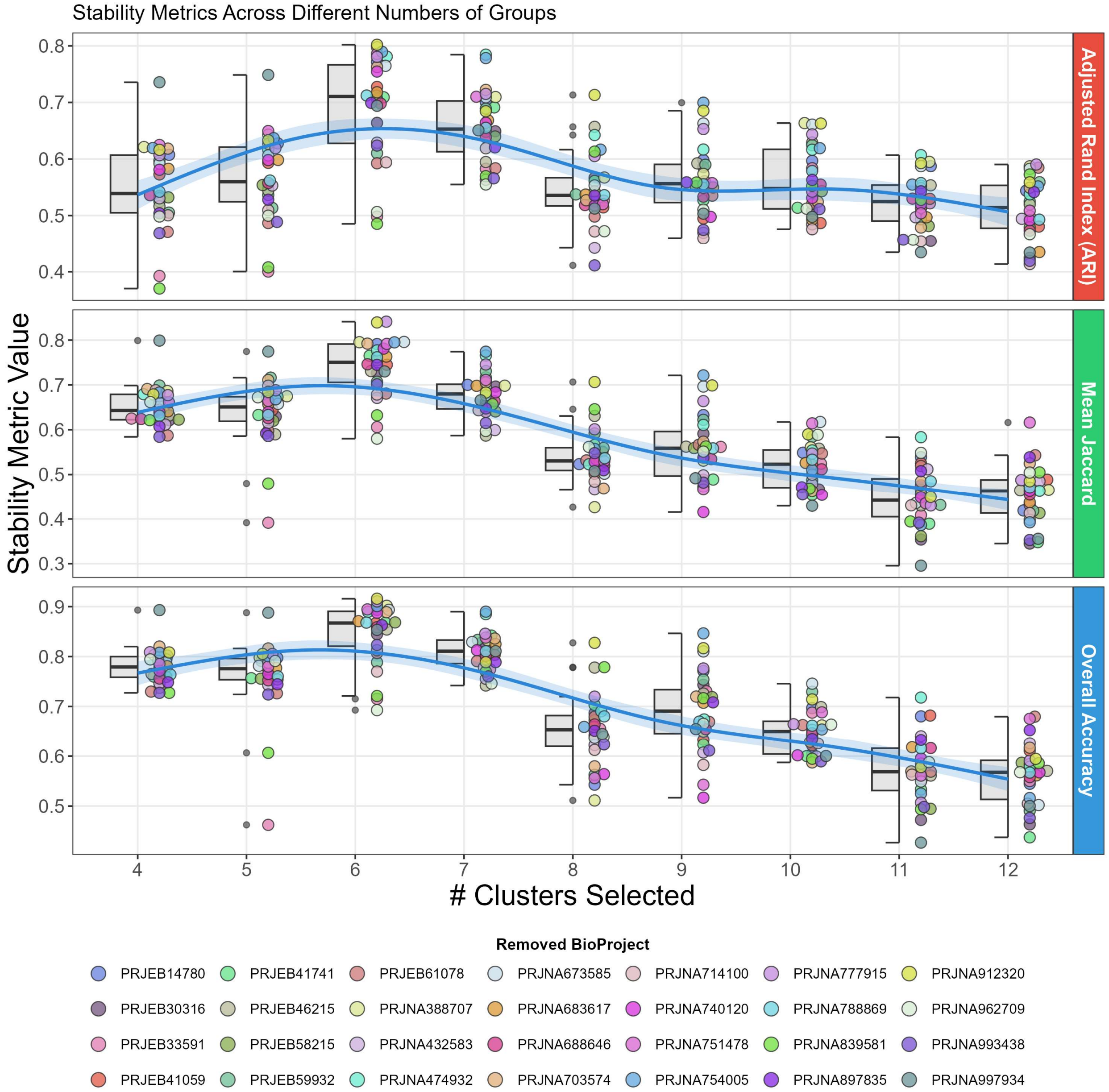


**Figure S5. Stability metrics across different cluster numbers evaluated through leave-one-study-out (LOSO) cross-validation**. Half boxplots (left, gray) show distribution quartiles for each cluster number. Individual data points (right, colored circles) represent stability measurements from the LOSO CV with different removed BioProjects. Blue trend lines show generalized additive model (GAM)-smoothed curves with 95% confidence intervals (shaded regions). Top panel: Adjusted Rand Index (ARI) measures clustering agreement corrected for chance. Middle panel: Mean Jaccard Index quantifies average cluster overlap. Bottom panel: Overall Accuracy represents proportion of correctly assigned samples.

#### Ulrb results

The ulrb analysis revealed distinct abundance patterns across all NPCSTs, with detailed results presented in **Figure S6-11**. The two diverse NPCSTs (V and VI) exhibited substantially higher numbers of abundant genera (94 and 65 genera, respectively) compared to the remaining NPCSTs (4-27 abundant genera each). This pattern corresponded well with the increased overall genera diversity observed in these two NPCSTs, confirming their classification as compositionally heterogeneous community types. One key distinguishing pattern among NPCSTs was the higher relative abundance of Streptococcus in specific community types. Silhouette score distributions consistently indicated strong clustering quality for genera abundance classifications across all NPCSTs, validating the robustness of the abundance categorizations.

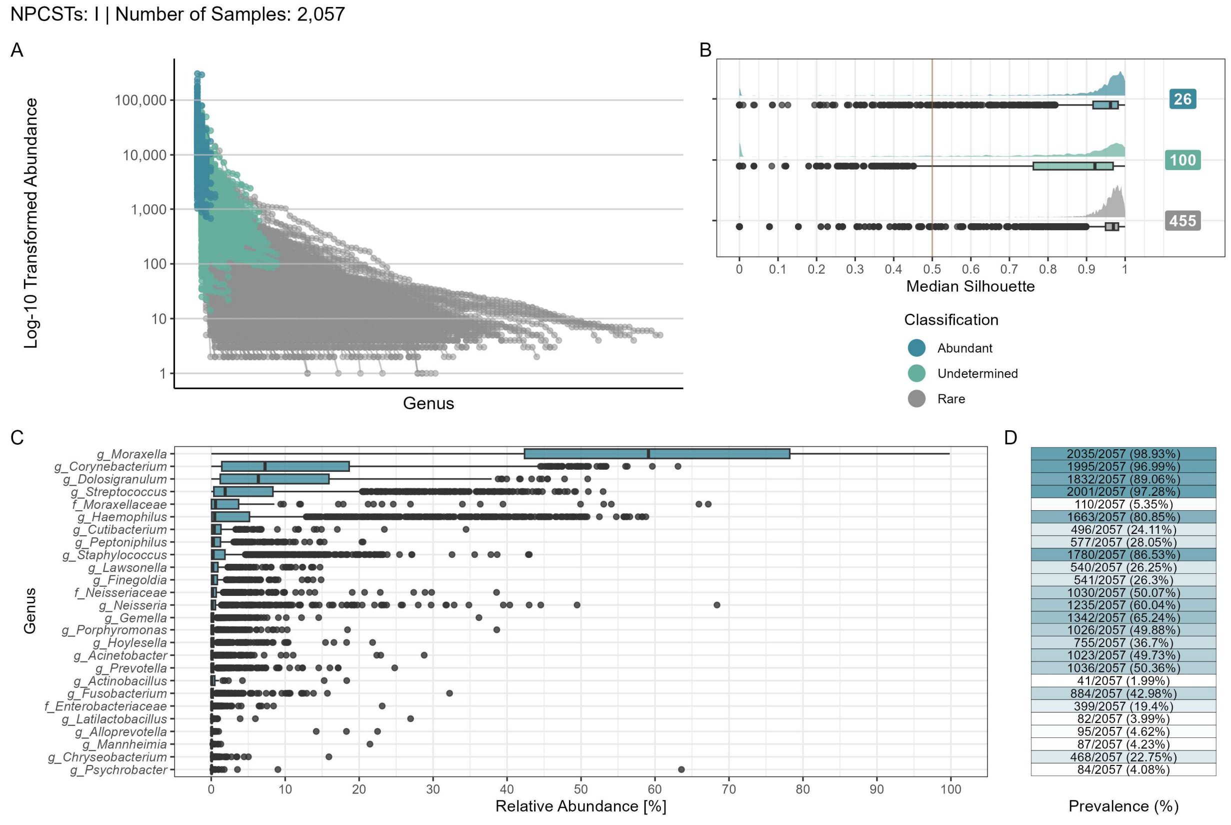


**Figure S6. Ulrb diagnostic plots for NPCST I (n = 2,057 samples). A.** $\text{Log}_{10}$-transformed relative abundance distribution across all genera, stratified by ulrb classifications ("Abundant", "Undetermined", and "Rare"). **B.** Silhouette score distributions shown as density plots and boxplots for each classification. The quality threshold (Silhouette score = 0.5) is indicated by the red dashed line. Numbers of unique genera per classification are displayed adjacent to each plot. **C.** Genus-level relative abundance for genera classified as "Abundant" by ulrb. **D.** Detection prevalence of abundant genera across samples, expressed as percentages. Classifications are color-coded: Abundant (blue), Undetermined (teal), and Rare (gray). Prevalence is represented on a gradient scale from 0-100% (white to blue).

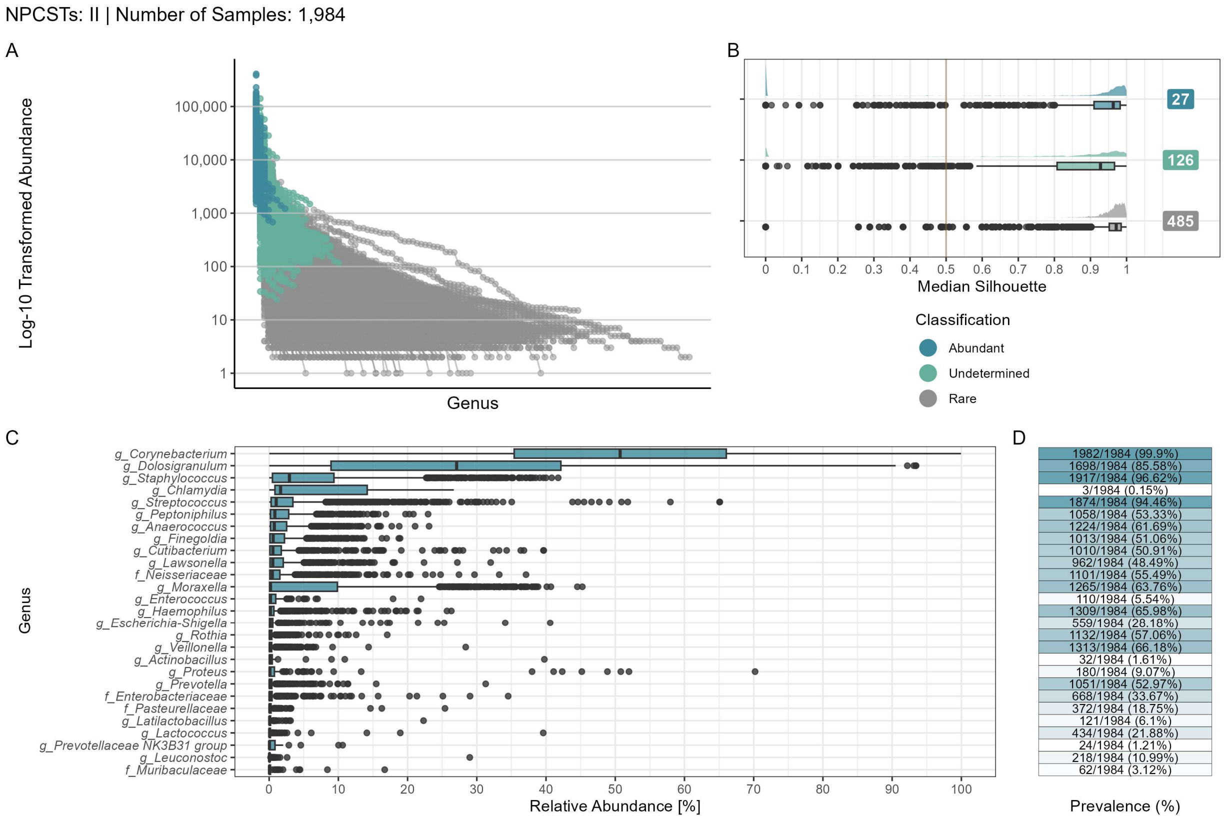


**Figure S7. Ulrb diagnostic plots for NPCST II (n = 1,984 samples). A.** Log$_{10}$-transformed relative abundance distribution across all genera, stratified by ulrb classifications ("Abundant", "Undetermined", and "Rare"). **B.** Silhouette score distributions shown as density plots and boxplots for each classification. The quality threshold (Silhouette score = 0.5) is indicated by the red dashed line. Numbers of unique genera per classification are displayed adjacent to each plot. **C.** Genus-level relative abundance for genera classified as "Abundant" by ulrb. **D.** Detection prevalence of abundant genera across samples, expressed as percentages. Classifications are color-coded: Abundant (blue), Undetermined (teal), and Rare (gray). Prevalence is represented on a gradient scale from 0-100% (white to blue).

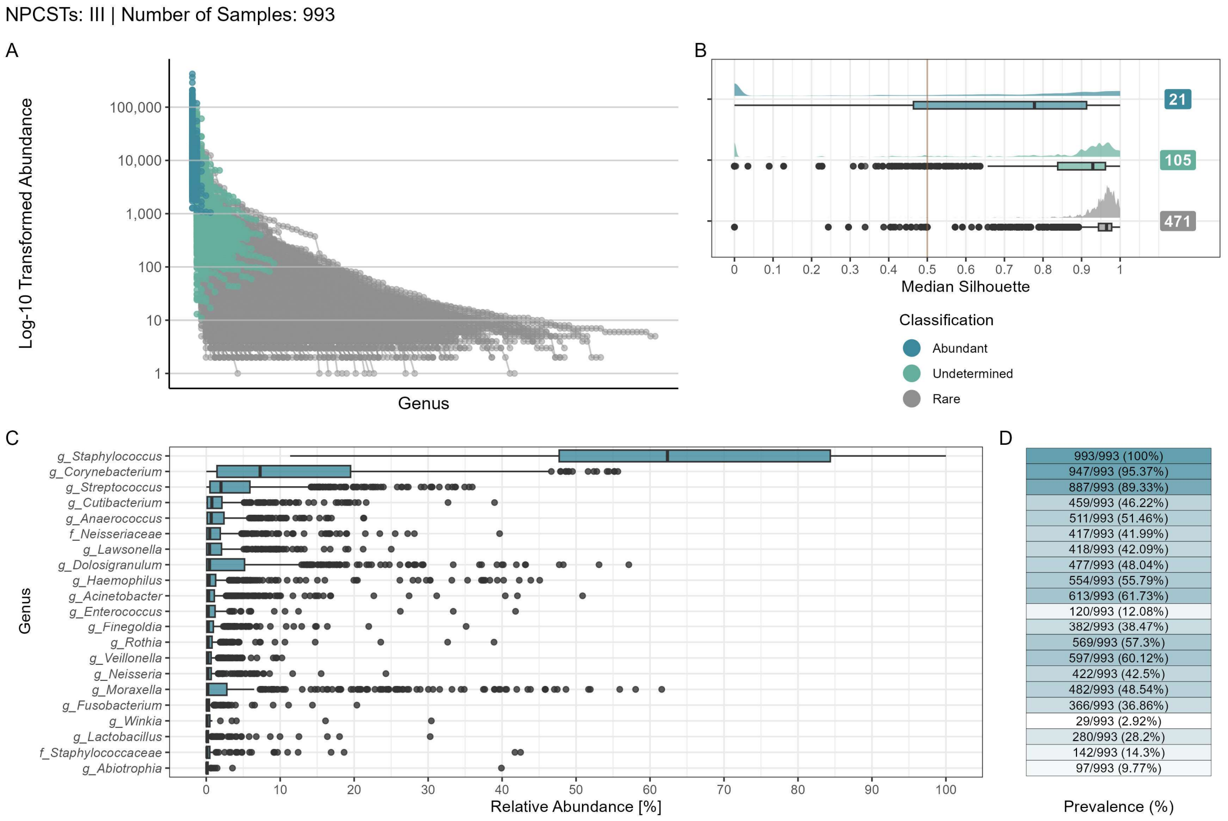


**Figure S8. Ulrb diagnostic plots for NPCST III (n = 993 samples). A.** Log$_{10}$-transformed relative abundance distribution across all genera, stratified by ulrb classifications ("Abundant", "Undetermined", and "Rare"). **B.** Silhouette score distributions shown as density plots and boxplots for each classification. The quality threshold (Silhouette score = 0.5) is indicated by the red dashed line. Numbers of unique genera per classification are displayed adjacent to each plot. **C.** Genus-level relative abundance for genera classified as "Abundant" by ulrb. **D.** Detection prevalence of abundant genera across samples, expressed as percentages. Classifications are color-coded: Abundant (blue), Undetermined (teal), and Rare (gray). Prevalence is represented on a gradient scale from 0-100% (white to blue).

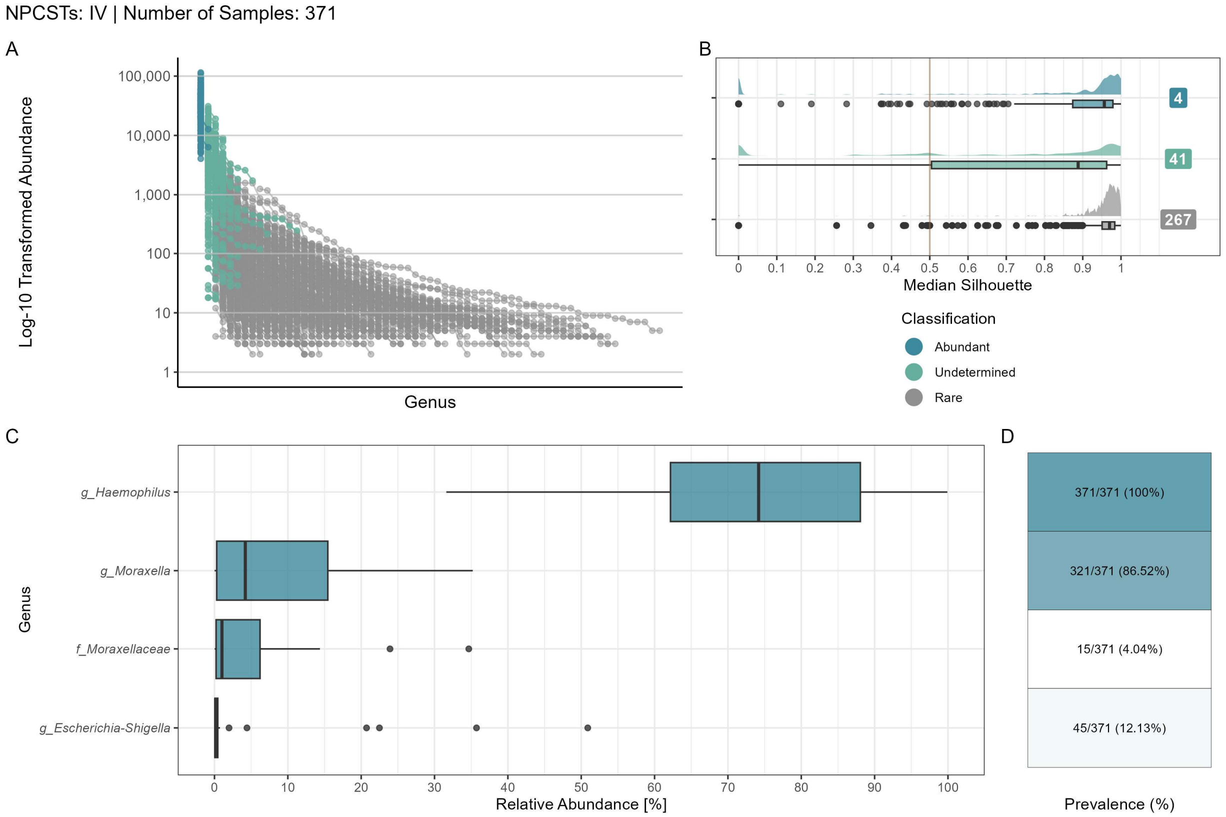


**Figure S9. Ulrb diagnostic plots for NPCST IV (n = 371 samples). A.** $Log_{10}$-transformed relative abundance distribution across all genera, stratified by ulrb classifications ("Abundant", "Undetermined", and "Rare"). **B.** Silhouette score distributions shown as density plots and boxplots for each classification. The quality threshold (Silhouette score = 0.5) is indicated by the red dashed line. Numbers of unique genera per classification are displayed adjacent to each plot. **C.** Genus-level relative abundance for genera classified as "Abundant" by ulrb. **D.** Detection prevalence of abundant genera across samples, expressed as percentages. Classifications are color-coded: Abundant (blue), Undetermined (teal), and Rare (gray). Prevalence is represented on a gradient scale from 0-100% (white to blue).

NPCSTs: V | Number of Samples: 961

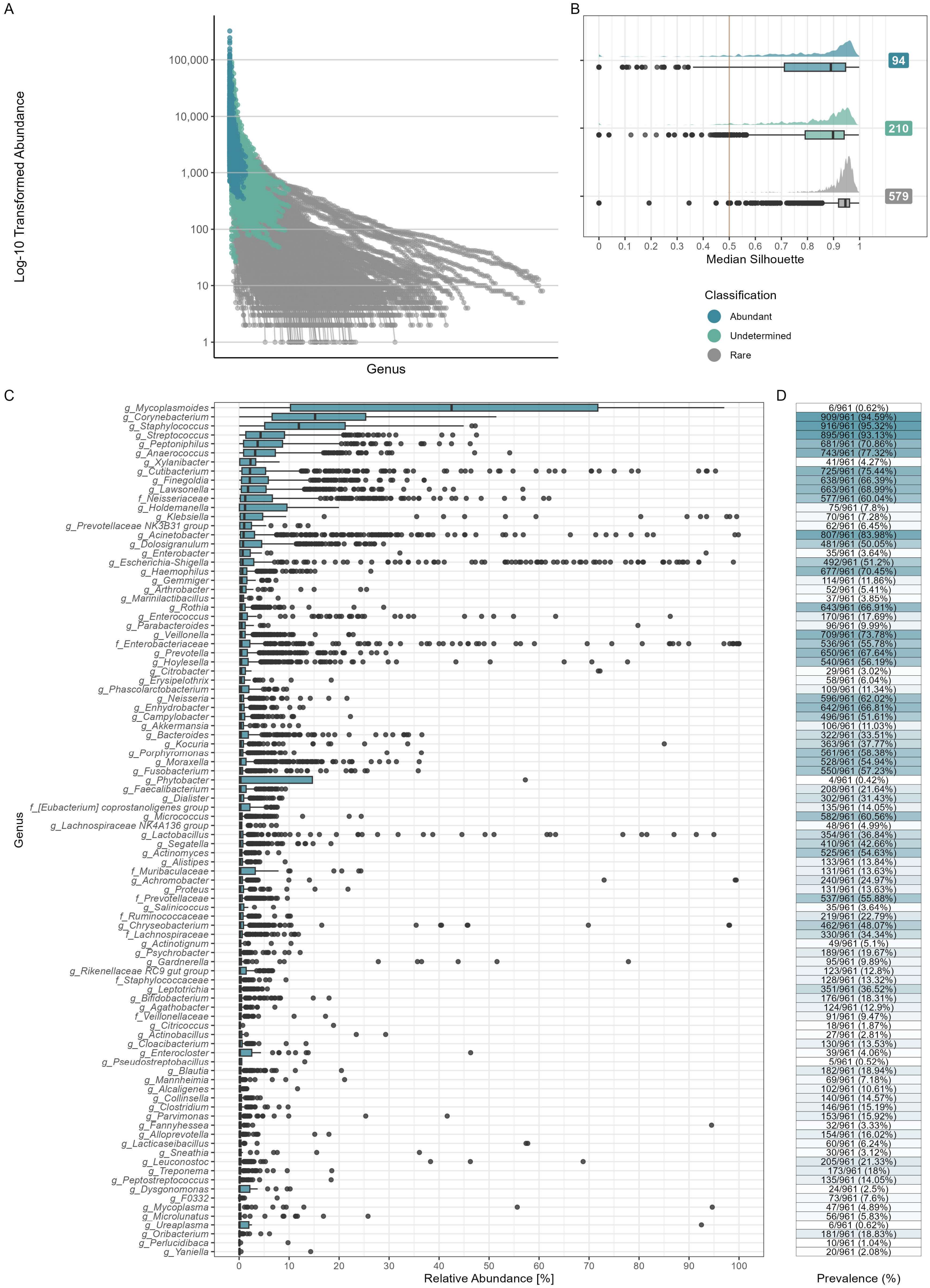

**Figure S10. Ulrb diagnostic plots for NPCST V (n = 961 samples). A.** $Log_{10}$-transformed relative abundance distribution across all genera, stratified by ulrb classifications ("Abundant", "Undetermined", and "Rare"). **B.** Silhouette score distributions shown as density plots and boxplots for each classification. The quality threshold (Silhouette score = 0.5) is indicated by the red dashed line. Numbers of unique genera per classification are displayed adjacent to each plot. **C.** Genus-level relative abundance for genera classified as "Abundant" by ulrb. **D.** Detection prevalence of abundant genera across samples, expressed as percentages. Classifications are color-coded: Abundant (blue), Undetermined (teal), and Rare (gray). Prevalence is represented on a gradient scale from 0-100% (white to blue).

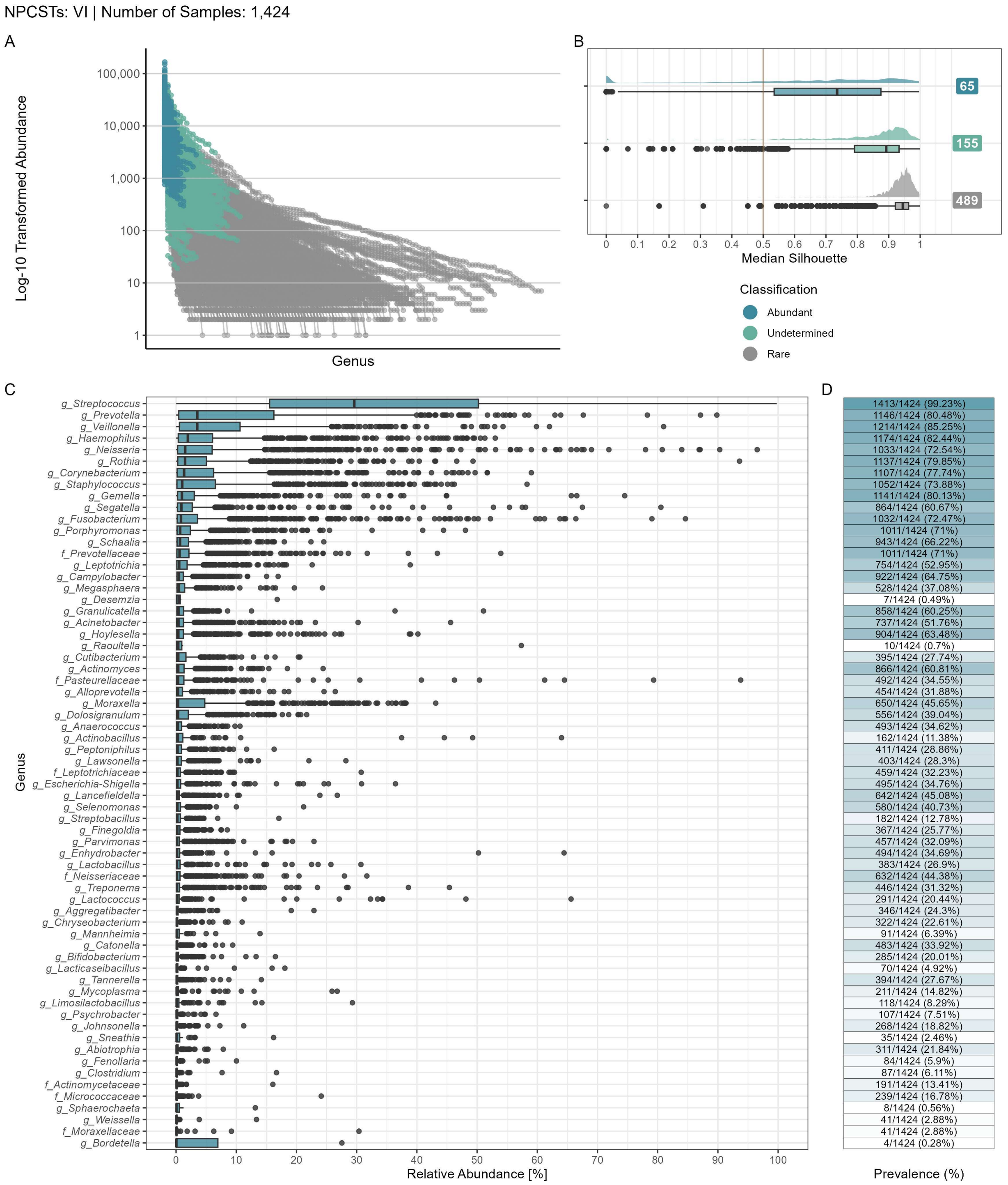


**Figure S11. Ulrb diagnostic plots for NPCST VI (n = 1,424 samples). A.** $Log_{10}$-transformed relative abundance distribution across all genera, stratified by ulrb classifications ("Abundant", "Undetermined", and "Rare"). **B.** Silhouette score distributions shown as density plots and boxplots for each classification. The quality threshold (Silhouette score = 0.5) is indicated by the red dashed line. Numbers of unique genera per classification are displayed adjacent to each plot. **C.** Genus-level relative abundance for genera classified as "Abundant" by ulrb. **D.** Detection prevalence of abundant genera across samples, expressed as percentages. Classifications are color-coded: Abundant (blue), Undetermined (teal), and Rare (gray). Prevalence is represented on a gradient scale from 0-100% (white to blue).

#### NPCST machine learning results

##### Model evaluation

Across 100 iterations of 5-fold cross-validation, machine learning models demonstrated strong performance for NPCST prediction. SVM achieved the highest performance (0.972), followed by Random Forest (0.966), Elastic Net (0.936), and Ridge (0.922). Given that Elastic Net and Ridge regression models showed significantly lower performance compared to SVM and Random Forest, we excluded these methods from detailed evaluation and focused on the two superior-performing algorithms. When examining per-NPCST performance, NPCST V (the diverse NPCST) demonstrated significantly reduced performance metrics compared to other NPCSTs. SVM achieved superior performance on NPCST V with 0.936 accuracy on the testing set, while Random Forest achieved 0.922.

For hyperparameter optimization, we evaluated 500 runs across 100 iterations of 5-fold cross-validation to identify optimal parameters. Random Forest hyperparameter optimization revealed mtry=209 (representing one-third of the 626 genera features) and ntree=50 as the most prevalent and optimal selection. Specifically, ntree=50 was optimal in 345/500 runs (69%), followed by ntree=100 in 152/500 runs (30.4%) and ntree=150 in 3/500 runs (0.6%), while mtry=209 was utilized across all runs. Importantly, our evaluation showed negligible performance differences between 50 and 100 trees, supporting selection of ntree=50 for computational efficiency. SVM optimization consistently identified cost=100 and gamma=1 as optimal parameters across all 500 runs (100% consistency).

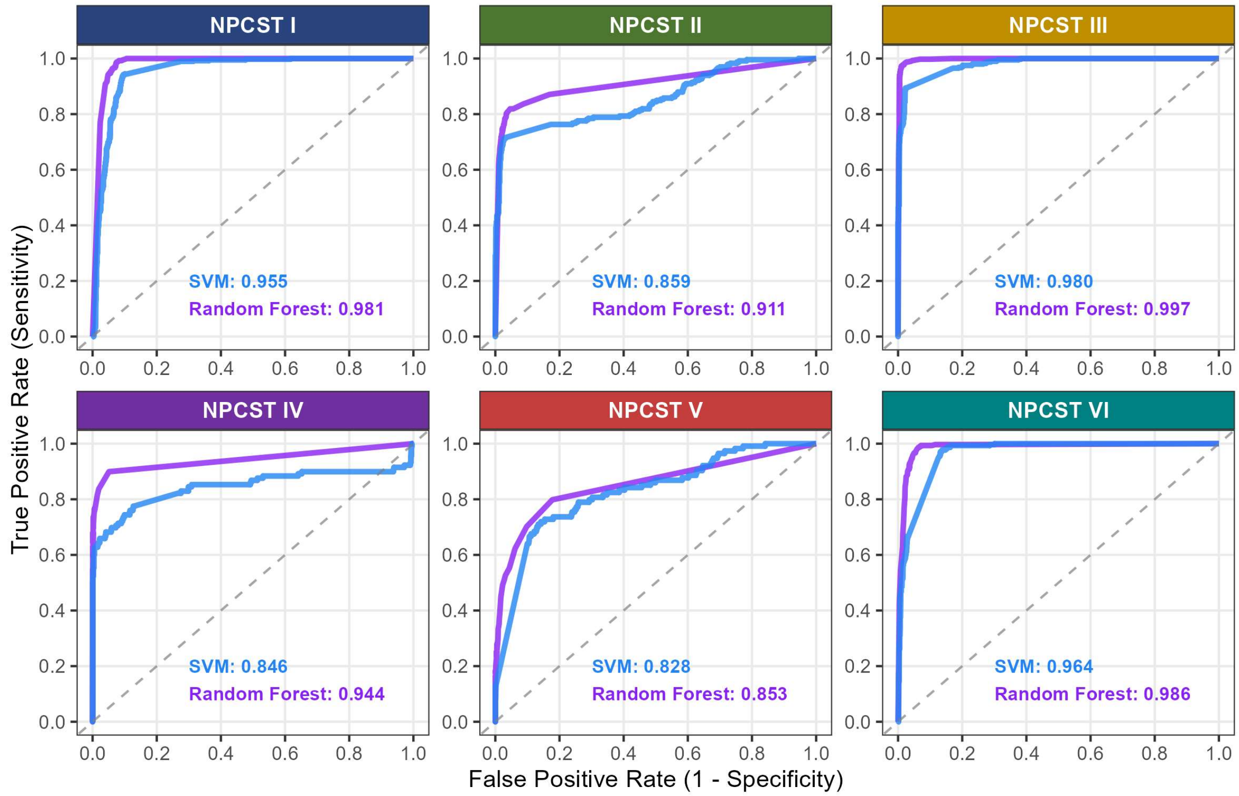


**Figure S12. ROC curve analysis of external validation NPCST predictions.** Ground truth assignments derived from hierarchical clustering analysis combining external validation studies with the original 28-study dataset (Bray-Curtis dissimilarity, Ward linkage; see **Methods** and **Supplementary File Methods**). Model performance is shown via ROC curves and AUC values for both SVM (blue) and Random Forest (purple) across all six NPCSTs.

#### Community-state-specific patterns in demographics, disease risk, and microbial diversity

NPCSTs demonstrated remarkable stability across demographic variables. Analysis of 5,353 cross-sectional samples yielded significant but small effect sizes: age ($\varepsilon^2 = 0.07$, 95% CI: 0.06–0.09) and sex (Cramér's V = 0.07, 95% CI: 0.04–0.09) each explained ≤7% of community variation (both $p < 0.0001$). The primary demographic pattern emerged in early childhood, with *Moraxella*-dominated (NPCST I) and *Haemophilus*-dominated (NPCST IV) communities preferentially colonizing children under 10 years (**Figure S13A-B**). The otherwise minimal demographic effects underscore the intrinsic stability of NPCSTs as fundamental ecological states of the nasopharyngeal microbiome.

We also observed significant variation in disease risk across NPCSTs when controlling for study effects (**Figure S13**). Most notably, bacterial infection risk was significantly higher in

NPCST VI communities (OR: 1.21; 95% CI: 1.10–1.33; $p < 0.012$) while viral infection risk was significantly lower (OR: 0.90; 95% CI: 0.86–0.94; $p < 0.001$). NPCST IV showed minimal but significant viral susceptibility (OR: 1.05; 95% CI: 1.00–1.10; $p = 0.028$). These findings implicate specific microbiome configurations in respiratory infection susceptibility.

Shannon diversity patterns were intrinsically linked to NPCST identity rather than disease status. Analysis of 5,000-read rarefied samples demonstrated three stable diversity categories: NPCST IV maintained consistently low diversity (<1.5), NPCSTs I–III showed intermediate levels (0–2.5), and NPCSTs V–VI exhibited high diversity (0–3.5) regardless of infection presence (**Figure S13D**). Disease-associated diversity changes within NPCSTs were minimal and inconsistent, while age contributed negligibly ($R^2 \leq 0.119$; **Figure S13E**). These stable, community-specific patterns have critical implications: diversity metrics must be interpreted within NPCST context.

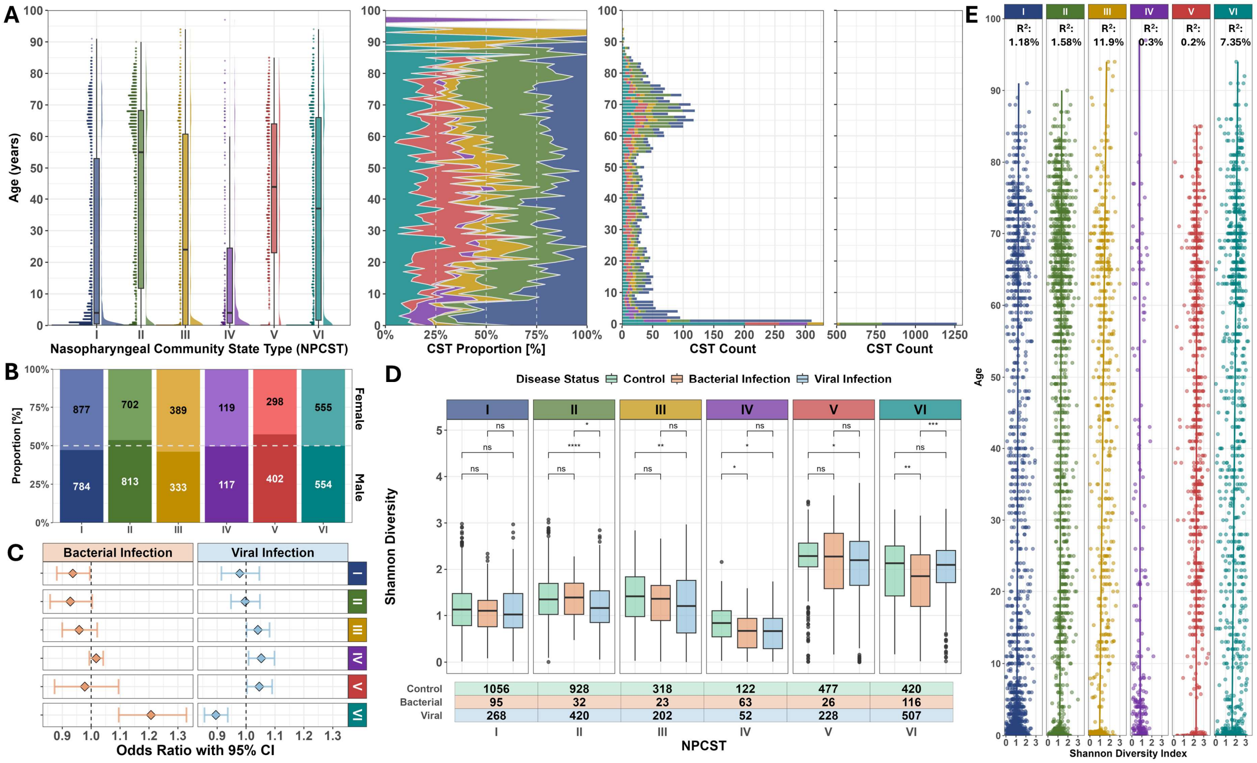


**Figure S13. Host demographic and ecological characteristics of nasopharyngeal community-state types (NPCSTs). A.** Age distribution across six NPCSTs displayed as raincloud plots combining density distributions (right), boxplots (center), and individual data points (left). The middle panel shows the proportion of each NPCST across age groups using 1-year increments. The two rightmost panels display the sample count for each NPCST at 1-year age increments. **B.** Sex distribution within each NPCST showing proportions of female (light shading) and male (dark shading) participants, with sample counts indicated within bars. Dashed white line indicates 50% proportion. **C.** Forest plot

displaying odds ratios (OR) with 95% confidence intervals for bacterial and viral infections compared to controls across NPCSTs, derived from multinomial logistic regression stratified by BioProject with OR=1 indicated by vertical black dashed lines. ORs >1.0 indicate increased infection risk, while ORs <1.0 indicate decreased risk relative to controls. **D** Shannon diversity distributions comparing control, bacterial infection, and viral infection groups within each NPCST. Sample sizes are shown below each group with significance levels from pairwise Wilcoxon rank-sum tests with FDR correction indicated above with brackets (*p < 0.05, **p < 0.01, ***p < 0.001, ns = not significant).
**E.** Relationship between Shannon diversity and age within each NPCST, with linear regression $R^2$ values displayed above each plot. Each point represents an individual sample colored by NPCST membership. In all panels, NPCSTs are color-coded as follows: I (blue), II (green), III (yellow), IV (purple), V (red), and VI (teal).

#### Co-occurrence network

**Figure S14A-B** presents centrality rankings (closeness, betweenness, and degree) for the top-performing genera. These 44 top-ranked genera (of 72 total analyzed) exhibited two distinct functional patterns: 17 multi-hub genera (including *g_Acinetobacter*, *g_Anaerococcus*, and *g_Peptoniphilus*) consistently ranked highly across all three network metrics in both NPCST-specific and global networks, indicating their role as keystone anchors that maintain community structure regardless of compositional shifts. The remaining 27 specialized genera showed prominence in only 1-2 metrics within specific NPCST contexts. For instance, *g_Enhydrobacter*, a gram-negative bacterium, ranked first for degree centrality in NPCST VI and first for closeness centrality in NPCSTs I and II, indicating its function as a context-specific influencer rather than a universal community hub. This specialization pattern provides new avenues for investigating these understudied genera. Importantly, *Moraxella* from NPCST I consistently showed low network centrality rankings despite its numerical dominance, demonstrating that competitive dominance can paradoxically result in network isolation and limited co-existence capacity. **Figure S14C** displays the co-occurrence networks for NPCSTs V and VI, which exhibit more complex association patterns consistent with their higher microbial diversity.

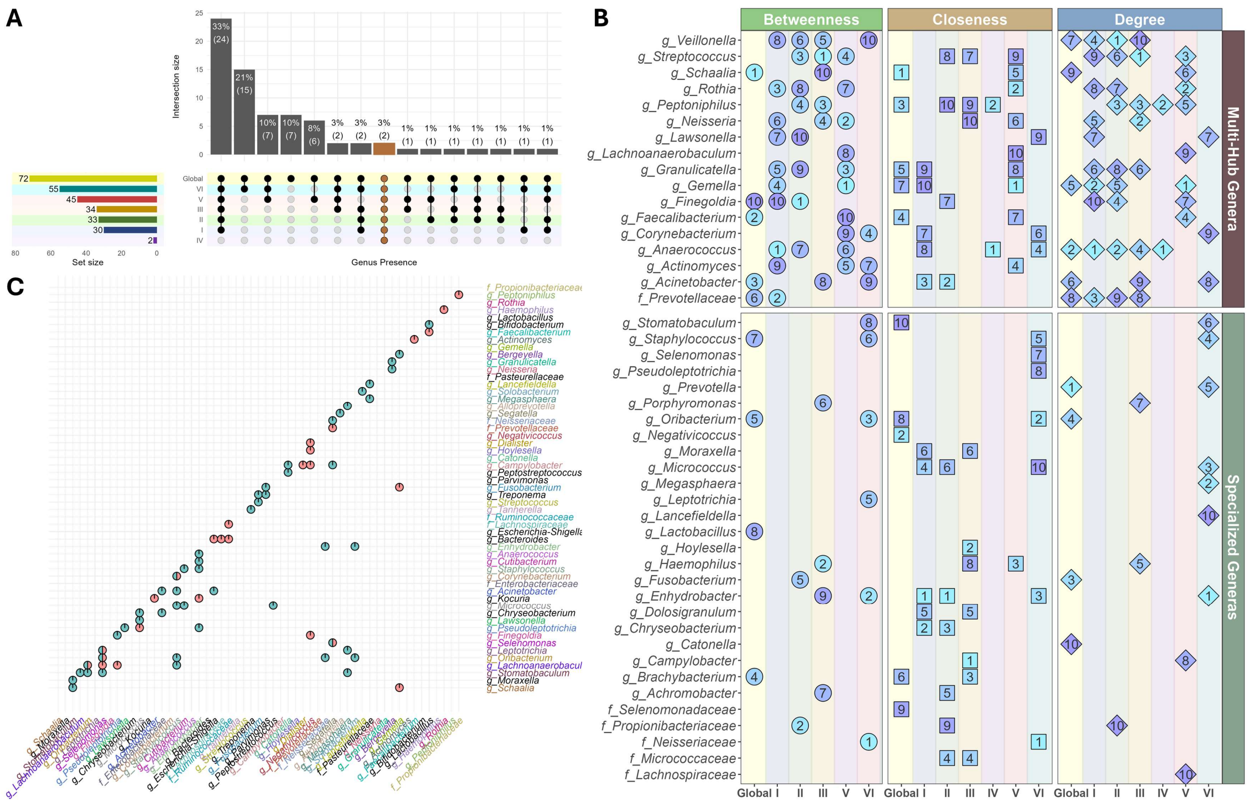


**Figure S14. Network Centrality Analysis of Structurally Important Genera. A.** UpSet plot showing the distribution of genera across global (orange dots and bar on upper marginal distribution) and NPCST-specific co-occurrence networks. Each column represents a unique combination of networks sharing specific genera, with the number and percentage of genera shown above each bar. The left panel indicates the total number of genera in each network (set size). The brown column highlights the, genera shared across all networks. Connected dots in the matrix indicate which networks contribute to each intersection. **B.** Centrality rankings (closeness, betweenness, and degree) for 44 top-ranked genera across the global and six NPCST-specific co-occurrence networks. Data point color indicates centrality ranking position (ranging from light blue=1 to purple=10), and shape indicates centrality type (circle=betweenness, square=closeness, and diamond=degree). Genera are stratified into two functional categories: multi-hub genera (n=17), representing taxa with consistently high centrality across multiple networks, and specialized genera (n=27), which exhibit prominent centrality within specific NPCST contexts but limited rankings across networks. **C.** Associations across networks, focusing on associations involving NPCSTs V and VI. Each pie chart represents microbial associations unique to NPCST V (red), unique to NPCST VI (teal), or shared between both NPCSTs (side by side color).

#### Co-detection of nasopharyngeal pathogens from PCR panels

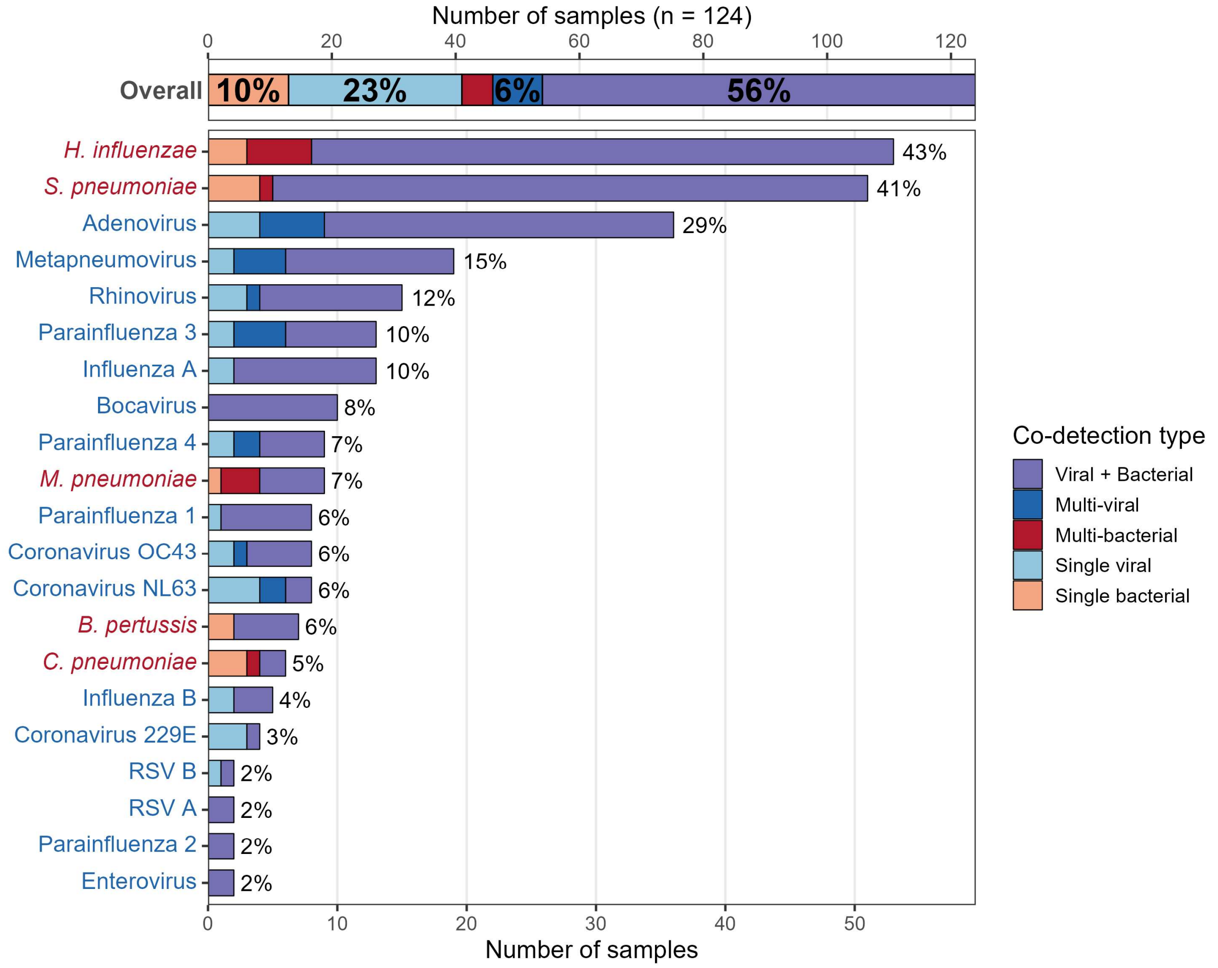


**Figure S15. Respiratory pathogen co-detection in nasopharyngeal clinical specimens. Multiplex RT-PCR (SeeGene NovaPlex™, 4 panels) applied to 124 remnant clinical nasopharyngeal swabs.** Top bar: overall distribution of co-detection categories across all specimens (n = 124 unique NP samples). Bottom bars: per-pathogen detection frequency with stacked segments indicating the co-detection context in which each pathogen was identified. Percentages denote the proportion of total specimens positive for each target. Bacterial species shown in red; viral targets in blue. Of all specimens, 66.9% harbored multiple pathogenic microbes simultaneously, demonstrating that poly-microbial colonization is the norm rather than the exception in the nasopharynx.

#### Nasopharyngeal microbiome health index external validation

With final model coefficients and optimal thresholds established, we evaluated NMHI performance on independent validation datasets excluded from all prior training stages. The external validation cohort was dominated by SARS-CoV-2 cases, reflecting continued research focus following the 2019 pandemic, with disease severity ranging from standard qPCR-confirmed SARS-CoV-2 to critical cases requiring ICU admission and mechanical ventilation. Additional validation samples included limited numbers of lower respiratory tract infections (LTRI, n=5), critically ill SARS-CoV-2-negative patients (n=31), and within the varying-severity SARS-CoV-2 group, symptomatic individuals with suspected but confirmed-negative SARS-CoV-2 (n=15). External validation maintained strong discriminatory performance, with Cohen's d values ranging from 1.598-2.368 across all diagnostic categories when compared to external healthy controls, indicating large effect sizes and robust separation between healthy and diseased populations. Notably, mechanically ventilated SARS-CoV-2 patients showed the lowest effect size, potentially due to procedural artifacts affecting the nasopharyngeal microbiome during or after intubation. Nevertheless, all disease categories demonstrated statistically significant distributional differences from healthy populations, confirming NMHI's broad applicability across diverse pathological conditions. AUC analysis further validated model performance, achieving 0.922 for the combined dataset and NPCST-specific values ranging from 0.848-0.953, with NPCST classifications determined using the random forest model developed in earlier stages to ensure reproducible workflow implementation. NPCST V was excluded from analysis as all samples belonged to the disease group, precluding meaningful AUC calculation.

### PRISMA checklist

| Section and Topic | Item # | Checklist item | Location where item is reported |
|---|---|---|---|
| **TITLE** | | | |
| Title | 1 | Identify the report as a systematic review. | Title |
| **ABSTRACT** | | | |
| Abstract | 2 | See the PRISMA 2020 for Abstracts checklist. | Abstract |
| **INTRODUCTION** | | | |
| Rationale | 3 | Describe the rationale for the review in the context of existing knowledge. | Introduction |
| Objectives | 4 | Provide an explicit statement of the objective(s) or question(s) the review addresses. | Introduction |
| **METHODS** | | | |
| Eligibility criteria | 5 | Specify the inclusion and exclusion criteria for the review and how studies were grouped for the syntheses. | Methods “Study screening and metadata evaluation“ & Figure 1B |
| Information sources | 6 | Specify all databases, registers, websites, organisations, reference lists and other sources searched or consulted to identify studies. Specify the date when each source was last searched or consulted. | Methods “Study screening and metadata evaluation“ |

| Section and Topic | Item # | Checklist item | Location where item is reported |
|---|---|---|---|
| Search strategy | 7 | Present the full search strategies for all databases, registers and websites, including any filters and limits used. | Methods "Study screening and metadata evaluation" |
| Selection process | 8 | Specify the methods used to decide whether a study met the inclusion criteria of the review, including how many reviewers screened each record and each report retrieved, whether they worked independently, and if applicable, details of automation tools used in the process. | Methods "Study screening and metadata evaluation" |
| Data collection process | 9 | Specify the methods used to collect data from reports, including how many reviewers collected data from each report, whether they worked independently, any processes for obtaining or confirming data from study investigators, and if applicable, details of automation tools used in the process. | Methods "16S data processing" |
| Data items | 10a | List and define all outcomes for which data were sought. Specify whether all results that were compatible with each outcome domain in each study were sought (e.g. for all measures, time points, analyses), and if not, the methods used to decide which results to collect. | Methods & Table 1 |
| | 10b | List and define all other variables for which data were sought (e.g. participant and intervention characteristics, funding sources). Describe any assumptions made about any missing or unclear information. | Table 1 |
| Study risk of bias assessment | 11 | Specify the methods used to assess risk of bias in the included studies, including details of the tool(s) used, how many reviewers assessed each study and whether they worked independently, and if applicable, details of automation tools used in the process. | Methods & Discussion (Limitations) |
| Effect measures | 12 | Specify for each outcome the effect measure(s) (e.g. risk ratio, mean difference) used in the synthesis or presentation of results. | Methods "Statistical Methods" |
| Synthesis methods | 13a | Describe the processes used to decide which studies were eligible for each synthesis (e.g. tabulating the study intervention characteristics and comparing against the planned groups for each synthesis (item #5)). | Methods "16S data processing" |
| | 13b | Describe any methods required to prepare the data for presentation or synthesis, such as handling of missing summary statistics, or data conversions. | Methods "16S data processing" & "Nasopharyngeal background decontamination protocol" |
| | 13c | Describe any methods used to tabulate or visually display results of individual studies and syntheses. | Methods (multiple sections) |
| | 13d | Describe any methods used to synthesize results and provide a rationale for the choice(s). If meta-analysis was performed, describe the model(s), method(s) to identify the presence and extent of statistical heterogeneity, and software package(s) used. | Methods (multiple sections) |
| | 13e | Describe any methods used to explore possible causes of heterogeneity among study results (e.g. subgroup analysis, meta-regression). | Methods (multiple sections) & Supplementary File |

| Section and Topic | Item # | Checklist item | Location where item is reported |
|---|---|---|---|
| | 13f | Describe any sensitivity analyses conducted to assess robustness of the synthesized results. | Methods (multiple sections) & Supplementary File |
| Reporting bias assessment | 14 | Describe any methods used to assess risk of bias due to missing results in a synthesis (arising from reporting biases). | Methods |
| Certainty assessment | 15 | Describe any methods used to assess certainty (or confidence) in the body of evidence for an outcome. | N/A |
| **RESULTS** | | | |
| Study selection | 16a | Describe the results of the search and selection process, from the number of records identified in the search to the number of studies included in the review, ideally using a flow diagram. | Methods “Study screening and metadata evaluation” |
| | 16b | Cite studies that might appear to meet the inclusion criteria, but which were excluded, and explain why they were excluded. | N/A |
| Study characteristics | 17 | Cite each included study and present its characteristics. | Table 1 |
| Risk of bias in studies | 18 | Present assessments of risk of bias for each included study. | N/A |
| Results of individual studies | 19 | For all outcomes, present, for each study: (a) summary statistics for each group (where appropriate) and (b) an effect estimate and its precision (e.g. confidence/credible interval), ideally using structured tables or plots. | N/A |
| Results of syntheses | 20a | For each synthesis, briefly summarise the characteristics and risk of bias among contributing studies. | Results (multiple sections) & Supplementary File |
| | 20b | Present results of all statistical syntheses conducted. If meta-analysis was done, present for each the summary estimate and its precision (e.g. confidence/credible interval) and measures of statistical heterogeneity. If comparing groups, describe the direction of the effect. | Results (multiple sections) & Supplementary File |
| | 20c | Present results of all investigations of possible causes of heterogeneity among study results. | Results (multiple sections) & Supplementary File |
| | 20d | Present results of all sensitivity analyses conducted to assess the robustness of the synthesized results. | Results (multiple sections) & Supplementary File |
| Reporting biases | 21 | Present assessments of risk of bias due to missing results (arising from reporting biases) for each synthesis assessed. | N/A |
| Certainty of evidence | 22 | Present assessments of certainty (or confidence) in the body of evidence for each outcome assessed. | Results (multiple sections on validations) |

| Section and Topic | Item # | Checklist item | Location where item is reported |
| --- | --- | --- | --- |
| | | | & Supplementary File on validations |
| **DISCUSSION** | | | |
| Discussion | 23a | Provide a general interpretation of the results in the context of other evidence. | Discussion |
| | 23b | Discuss any limitations of the evidence included in the review. | Discussion |
| | 23c | Discuss any limitations of the review processes used. | Discussion |
| | 23d | Discuss implications of the results for practice, policy, and future research. | Discussion |
| **OTHER INFORMATION** | | | |
| Registration and protocol | 24a | Provide registration information for the review, including register name and registration number, or state that the review was not registered. | N/A |
| | 24b | Indicate where the review protocol can be accessed, or state that a protocol was not prepared. | N/A |
| | 24c | Describe and explain any amendments to information provided at registration or in the protocol. | N/A |
| Support | 25 | Describe sources of financial or non-financial support for the review, and the role of the funders or sponsors in the review. | Funding statement |
| Competing interests | 26 | Declare any competing interests of review authors. | Conflict of Interest |
| Availability of data, code and other materials | 27 | Report which of the following are publicly available and where they can be found: template data collection forms; data extracted from included studies; data used for all analyses; analytic code; any other materials used in the review. | Data availability |

### Reference

Automatic citation updates are disabled. To see the bibliography, click Refresh in the Zotero tab.
